# Accelerated EM Connectome Reconstruction using 3D Visualization and Segmentation Graphs

**DOI:** 10.1101/2020.01.17.909572

**Authors:** Philip M. Hubbard, Stuart Berg, Ting Zhao, Donald J. Olbris, Lowell Umayam, Jeremy Maitin-Shepard, Michal Januszewski, William T. Katz, Erika R. Neace, Stephen M. Plaza

## Abstract

Recent advances in automatic image segmentation and synapse prediction in electron microscopy (EM) datasets of the brain enable more efficient reconstruction of neural connectivity. In these datasets, a single neuron can span thousands of images containing complex tree-like arbors with thousands of synapses. While image segmentation algorithms excel within narrow fields of views, the algorithms sometimes struggle to correctly segment large neurons, which require large context given their size and complexity. Conversely, humans are comparatively good at reasoning with large objects. In this paper, we introduce several semi-automated strategies that combine 3D visualization and machine guidance to accelerate connectome reconstruction. In particular, we introduce a strategy to quickly correct a segmentation through merging and *cleaving*, or splitting a segment along supervoxel boundaries, with both operations driven by affinity scores in the underlying segmentation. We deploy these algorithms as streamlined workflows in a tool called *Neu3* and demonstrate superior performance compared to prior work, thus enabling efficient reconstruction of much larger datasets. The insights into proofreading from our work clarify the trade-offs to consider when tuning the parameters of image segmentation algorithms.

## 1 Introduction

EM connectomics is an emerging field where the goal is to extract neural connectivity from high-resolution EM datasets. Since synapses and fine neuronal processes are very small, the datasets require nanometer-level resolution meaning that even a small organism requires several terabytes of image data [1]. These dataset sizes pose scalability challenges to both image acquisition and analysis. Recent advances in image segmentation [2, 3] and synapse prediction [4] make it possible to extract connectivity information data much faster than manually annotating the raw data [5].

Despite these advances, extensive manual validation of these automated predictions is still needed. For instance, tools like NeuTu [6] and webKnossis [7] allow proofreaders to manually merge or split segments to correct image segmentation errors. While these tools provide powerful editing capabilities, they often require humans to scroll through extensive image data to make time-consuming low-level decisions about errors in segmentation. Furthermore, splitting a falsely merged neuron is often very complicated and time consuming. The example in Figure 1 shows a segment that falsely merges two neuron fragments. Identifying the precise sources of merge errors and untangling these neurons is difficult. Additionally, the algorithms for semi-automated splitting [6], though comprehensive, are often time consuming to run for big neurons. As a result, segmentation algorithms are often tuned to conservatively avoid these type of false mergers, at the detriment of introducing false split errors.

**Figure 1:**
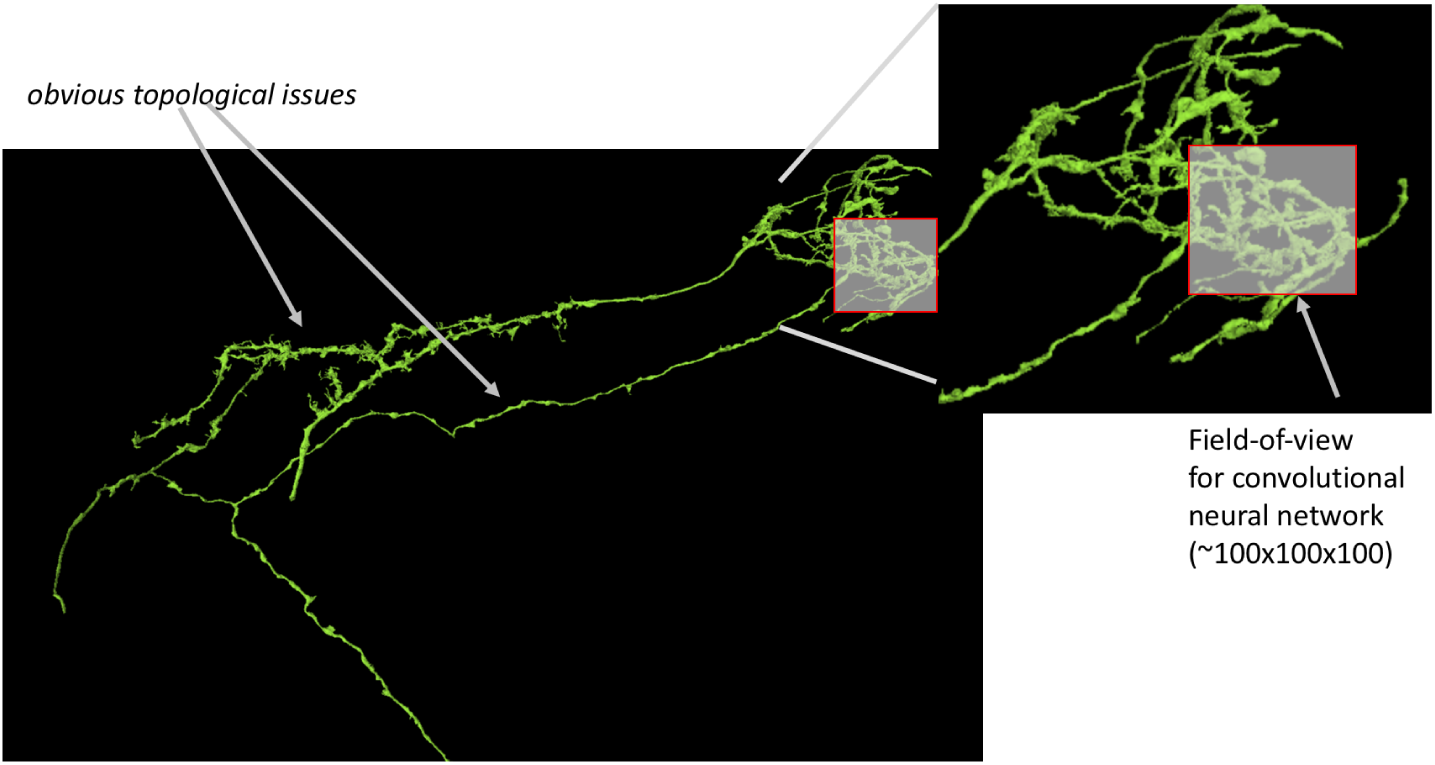
Segmentation algorithms perform well in a small local context as they have a limited field of view. Small local errors can result in large topological errors that are readily identifiable by humans. Reprinted from “Analyzing Image Segmentation for Connectomics,” by S. Plaza and J. Funke, 2018, *Frontiers in Neural Circuits*(18). Copyright 2018, Plaza and Funke. Reprinted with permission.

Our work is based on the following observations:

1. Current image segmentation often contains segments correct over spans of hundreds of image planes (i.e., several microns) consisting of large fragments of a neuron’s shape
2. Segmentation typically uses local context and therefore generally makes accurate predictions at the local level. However, subtle errors often manifest into large errors in the high-dimensional space as seen in Figure 1.
3. Humans’ visual processing abilities often enable them to identify subtle differences in 3D structures, such as tree-like branching patterns, as evidence in a study where around 60,000 trees were manually cataloged [8]. We expect humans to identify large errors in neuron shape with ease.
4. Many segmentation approaches start from an initial conservative base segmentation into *supervoxels* and involve a series of agglomeration steps to produce a more aggressive segmentation.

We exploit these observations in a new tool, *Neu3*, that provides a top-down 3D visualization environment to enable rapid examination of large segments and a fast, semi-automated re-segmentation strategy to quickly *cleave* falsely merged segments at supervoxel boundaries. With this approach, one can start with a more aggressive initial segmentation and spend time proofreading, at the object level, the large segments in the volume (which correlate in number with the neurons intersecting the dataset). In particular, we introduce the following:

1. 3D mesh visualization that highlights underlying supervoxels, which helps to visually decompose complex neuron shapes, along with a bird’s-eye view to provide multiple levels of 3D viewing.
2. The ability to rapidly view large bodies with *hybrid visualization*, combining high resolution at the center of attention and low resolution elsewhere.
3. A fast graph-cut re-segmentation algorithm for cleaving a segment along its supervoxel boundaries, implemented in conjunction with DVID [9], a web service that supports data versioning.
4. A 3D supervoxel splitting algorithm for the less common case of false merges within supervoxels.
5. A 3D-oriented form of focused proofreading [10] we call *focused merging*, based on agglomeration confidence, for fixing false splits at supervoxel boundaries.
6. Several proofreading protocols in Neu3 to streamline proofreading tasks.

We demonstrate the effectiveness of these contributions in Neu3 on a large, unpublished dataset, where we show that cleaving offers interactive performance that scales well, and that focused merging maintains high throughput. With these improvements, it is possible to rapidly review neurons at a high-level making the proofreading of even larger organisms possible.

The paper is organized as follows. After an overview of the proofreading process, we introduce 3D visualization enhancements in our proofreading workflows, and the semi-automated merging and cleaving strategies that leverage these enhancements. Then we describe different streamlined proofreading workflows implemented in Neu3. Finally, we present results and conclusions.

## 2 Proofreading Overview

Our proofreading process is based on the idea of a *protocol*, in which a proofreader performs instances of the same well-defined task. Protocols aim to minimize the choices a proofreader must make in each task, thus making the proofreading process more streamlined and efficient.

For the cleaving protocol, each task presents the proofreader with a set of supervoxels thought to define a single agglomerated body. If the proofreader believes more than one body is present, the proofreader designates supervoxels belonging to distinct agglomerated bodies and lets a server partition all the supervoxels accordingly, as described in Section 4. This approach is efficient because the proofreader chooses among existing supervoxels, and is freed from drawing free-form boundaries for splitting. For the focused merging protocol, each task presents the proofreader with two bodies, each containing one supervoxel from a pair with a strong agglomeration affinity. The proofreader determines if the two bodies should be merged into one. This approach is efficient because the proofreader is making only a binary decision. More details of the protocol workflows appear in Section 5.

Cleaving and focused merging involve merely redefining how supervoxels are grouped into bodies. The supervoxels themselves are immutable, which improves the performance of the system. For example, some of the visualization techniques described in Section 3 involve relatively high-resolution meshes of the supervoxels’ surfaces, and since the supervoxels are immutable, these meshes can be precomputed before proofreading begins.

Our supervoxels are large enough that a small fraction contain false merges, which cannot be fixed by cleaving. Therefore, the user interface for the cleaving protocol includes optional controls for free-form splitting of a supervoxel. The protocol allows the proofreader to proceed with supervoxel splitting as the need is encountered, or to mark it to be done later, to preserve the flow of the protocol. Supervoxel splitting is a rare situation that requires the regeneration of some meshes.

Data like supervoxel meshes and associations between supervoxels and agglomerated bodies are stored in the DVID system [9]. DVID supports branched versioning to maintain data provenance during proofreading, and provides a REST API that translates high-level operations like cleaving and merging to low-level updates to an underlying key-value store. Associations between supervoxels and bodies are maintained by DVID’s *labelmap* data type. The labelmap type employs a versioned map of labels to represent how atomic supervoxels are agglomerated into larger bodies, also designated by labels. Cleaving and merging are efficient because this map is cached in memory, and runtime is proportional to the number of supervoxels in a body rather than the number of voxels in the dataset. Supervoxel splitting operations are not as efficient but they are significantly less common in our workflows.

## 3 3D Visualization

Our 3D visualization for cleaving uses precomputed supervoxel meshes, since supervoxels are immutable for most proofreading operations as described in the previous section. DVID has a data type to store all the supervoxel meshes for an agglomerated body as a tar archive. Each mesh is compressed in Draco format [11] to reduce transmission time. Neu3 further improves performance by decompressing the meshes from an archive in parallel, and by prefetching the meshes for the next task. All the supervoxel meshes from one body can be displayed in the same color, or each can be colored independently with colors of random hue. Proofreaders find that coloring supervoxels independently helps them make sense of bodies with complex morphology. To further handle complexity, a proofreader can select an arbitrary set of supervoxel meshes and hide them to declutter the visualization. Such hiding is especially useful for supervoxel splitting, which requires the user to focus on the details of a single supervoxel mesh.

Our proofreaders have found it helpful to use a relatively high resolution for all supervoxel meshes when cleaving, since it may be necessary to look anywhere to find where a body should be cleaved. In focused merging, on the other hand, there is a predicted merge position for the two bodies in each task. As described in Section 4.2, this merge position lies midway on a line segment, *L*, between a point on each body. We exploit this information with a *hybrid visualization* mode, in which each body is displayed at a low resolution except in a small region of interest (ROI) around the merge position, where a high resolution is used. The ROI is an axis-aligned box, sized so that the rendering camera’s view direction is large enough to bound both bodies (Figure 2). The meshes used in this mode are not precomputed, but generated dynamically by Neu3 as it presents the task to the proofreader. There is no stitching on the boundary between the meshes inside and outside the ROI; small gaps thus may appear on that boundary, as shown in Figure 13, but they do not cause problems in practice. Dynamic generation of the meshes is efficient if the ROI is reasonably small, and the runtime compares favorably with the time required to download and decompress the precomputed supervoxel meshes as the pairs of bodies get larger.

**Figure 2:**
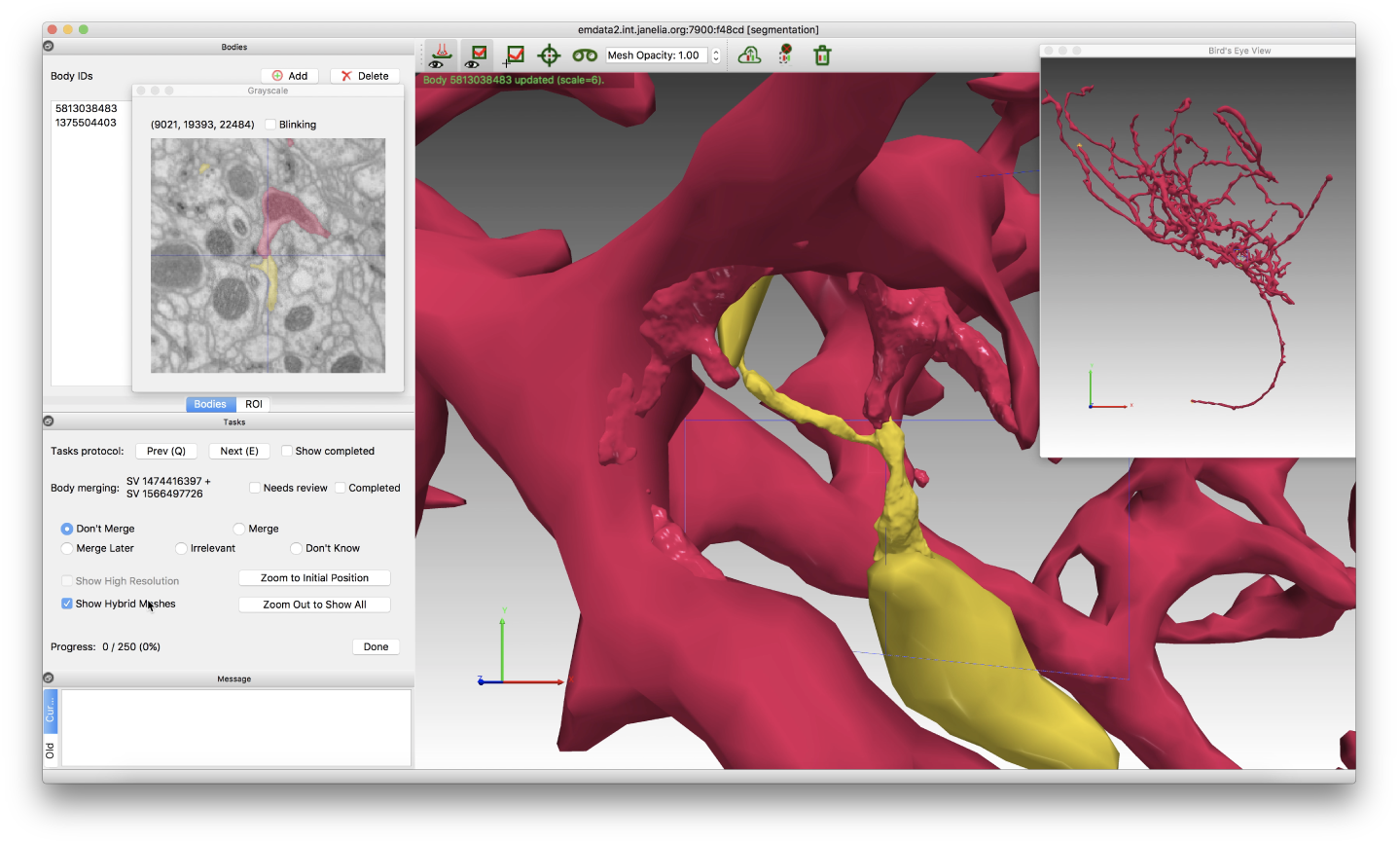
The ROI for this hybrid visualization bounds both bodies in the *z* dimension, the dimension most aligned with the camera’s view direction.

**Figure 3:**
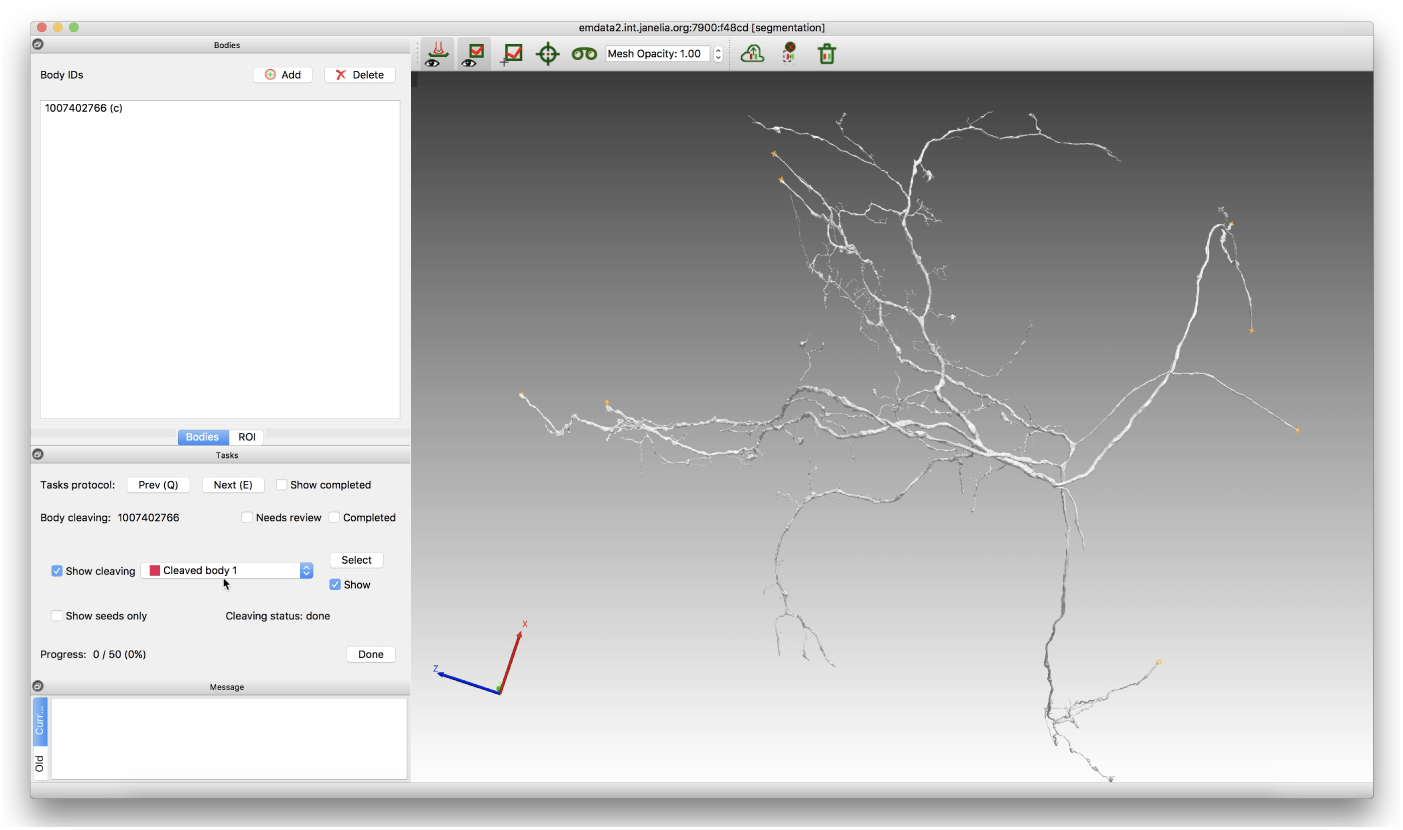
The Neu3 user interface at the start of a cleaving task for an agglomerated body with a false merge.

**Figure 4:**
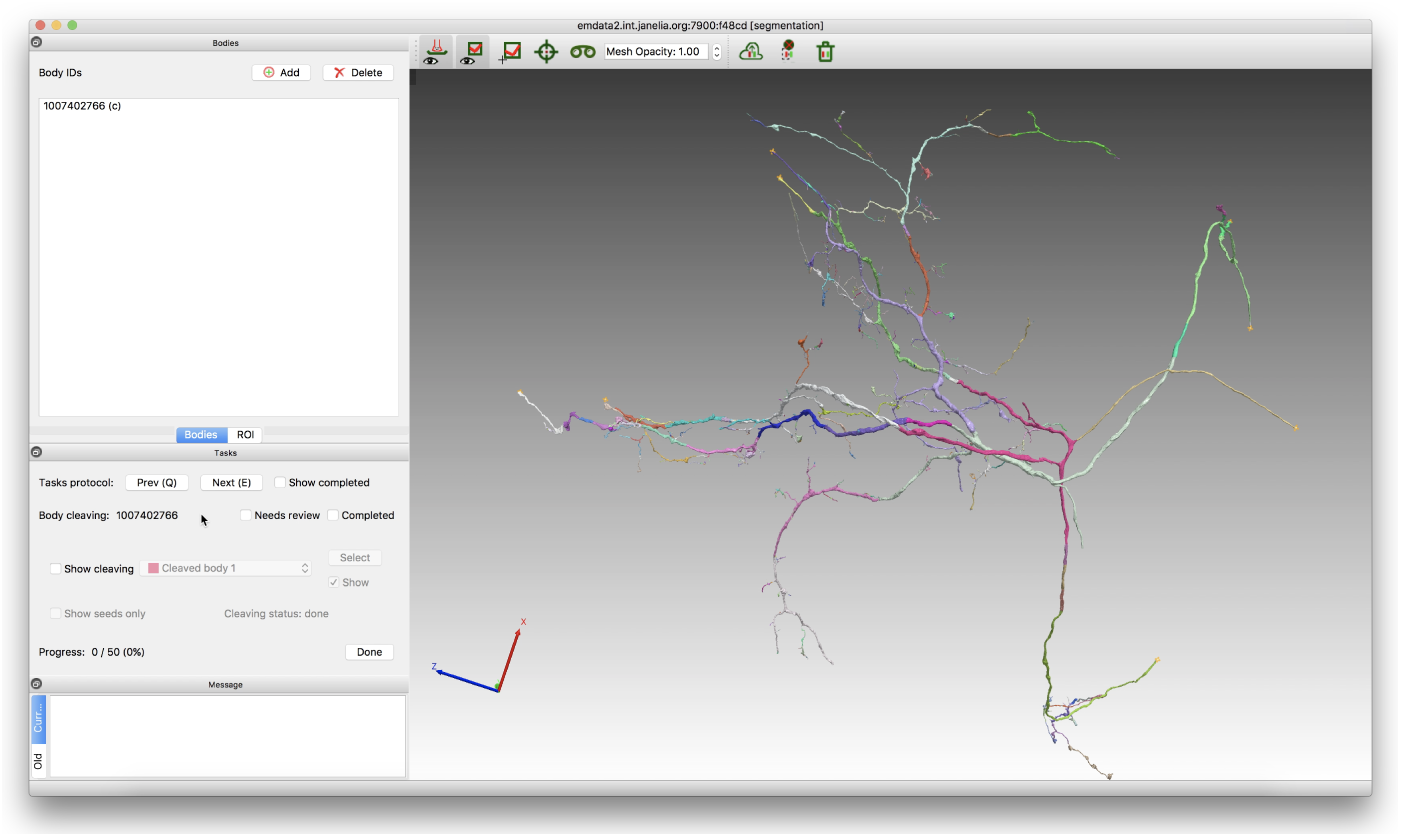
The Neu3 user interface for cleaving, with supervoxels colored independently.

**Figure 5:**
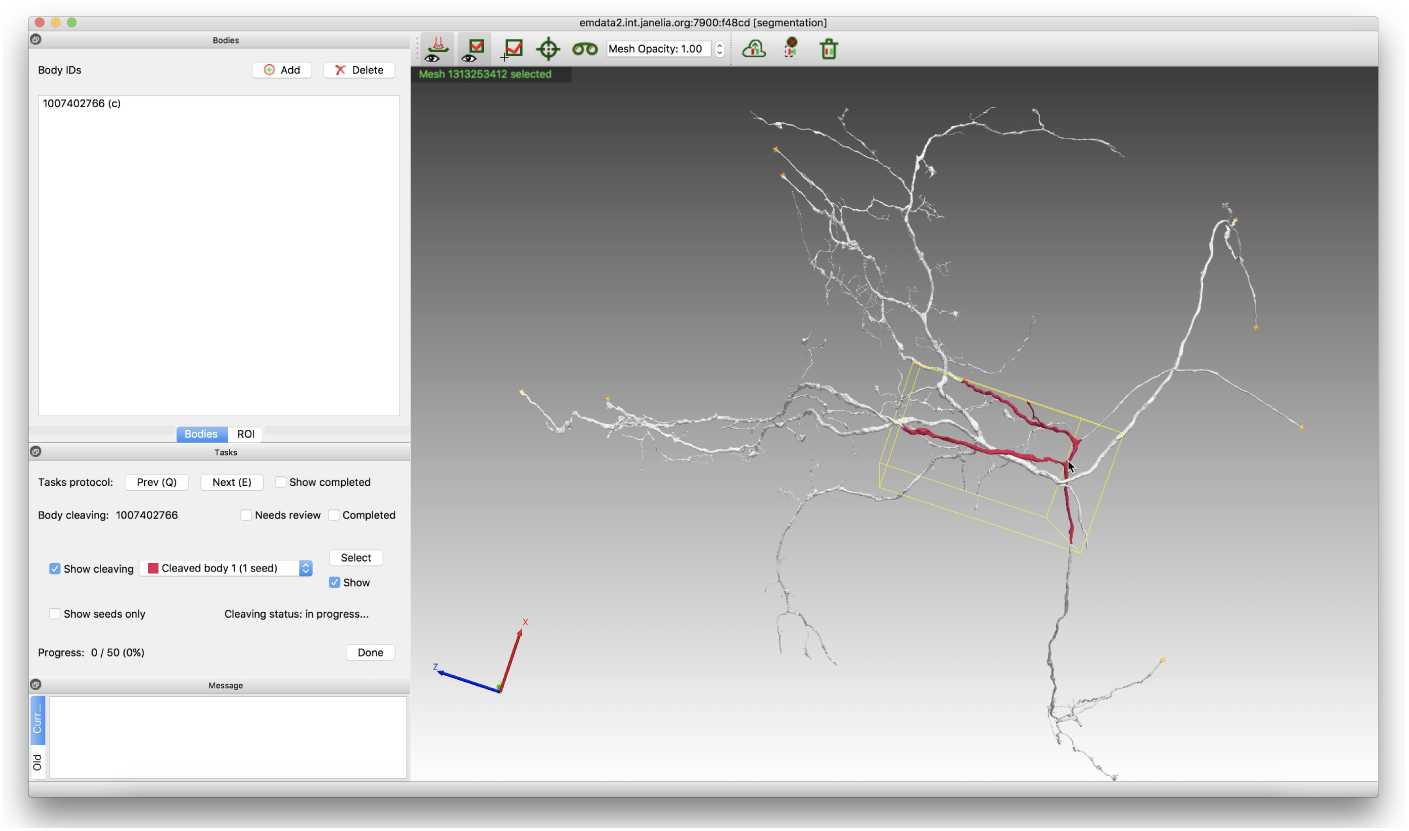
The Neu3 user interface for cleaving, showing a seed supervoxel for the first body. The yellow box indicates that the mesh is selected.

**Figure 6:**
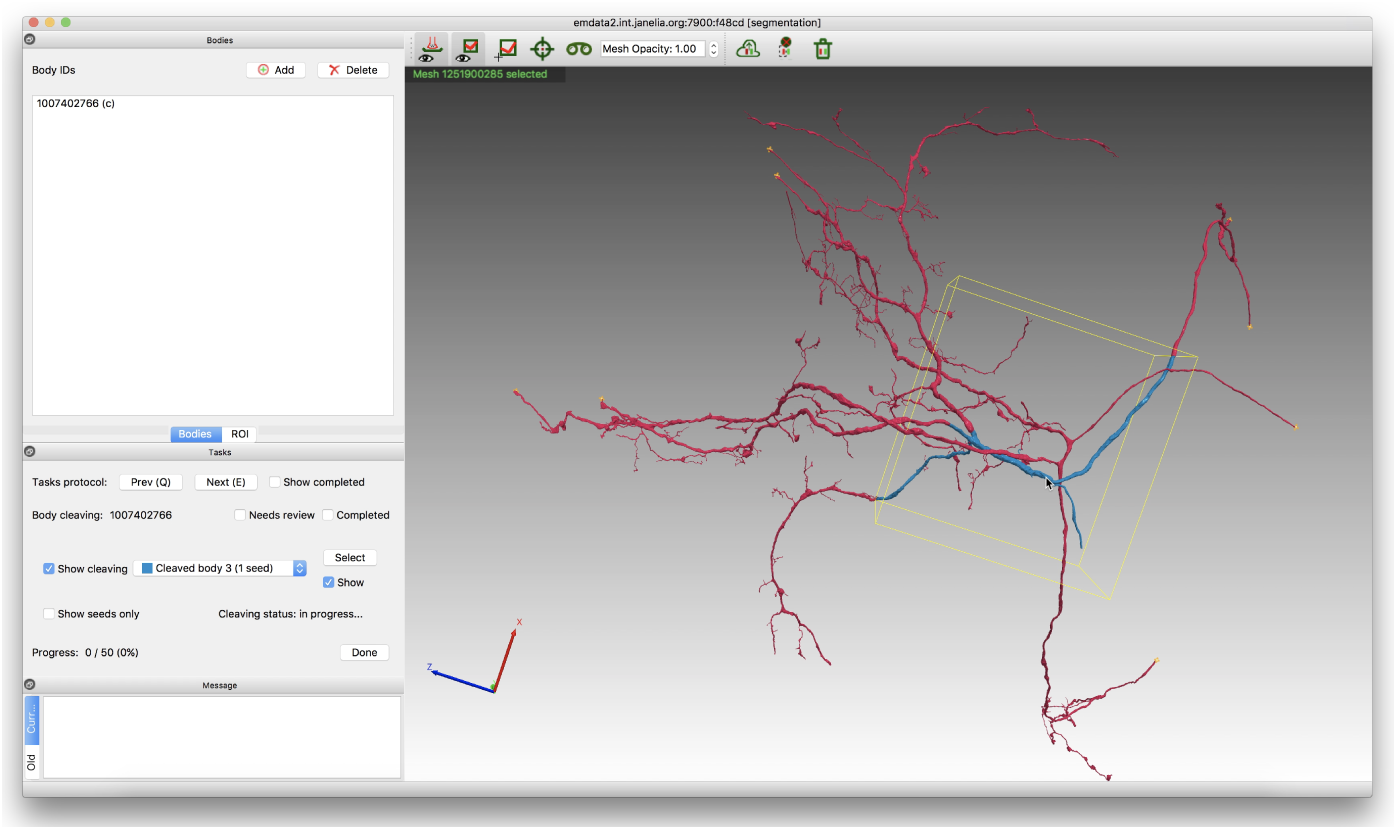
The Neu3 user interface for cleaving, showing a seed supervoxel for the second body. Until this seed is processed, everything else is considered part of the first body and colored accordingly.

**Figure 7:**
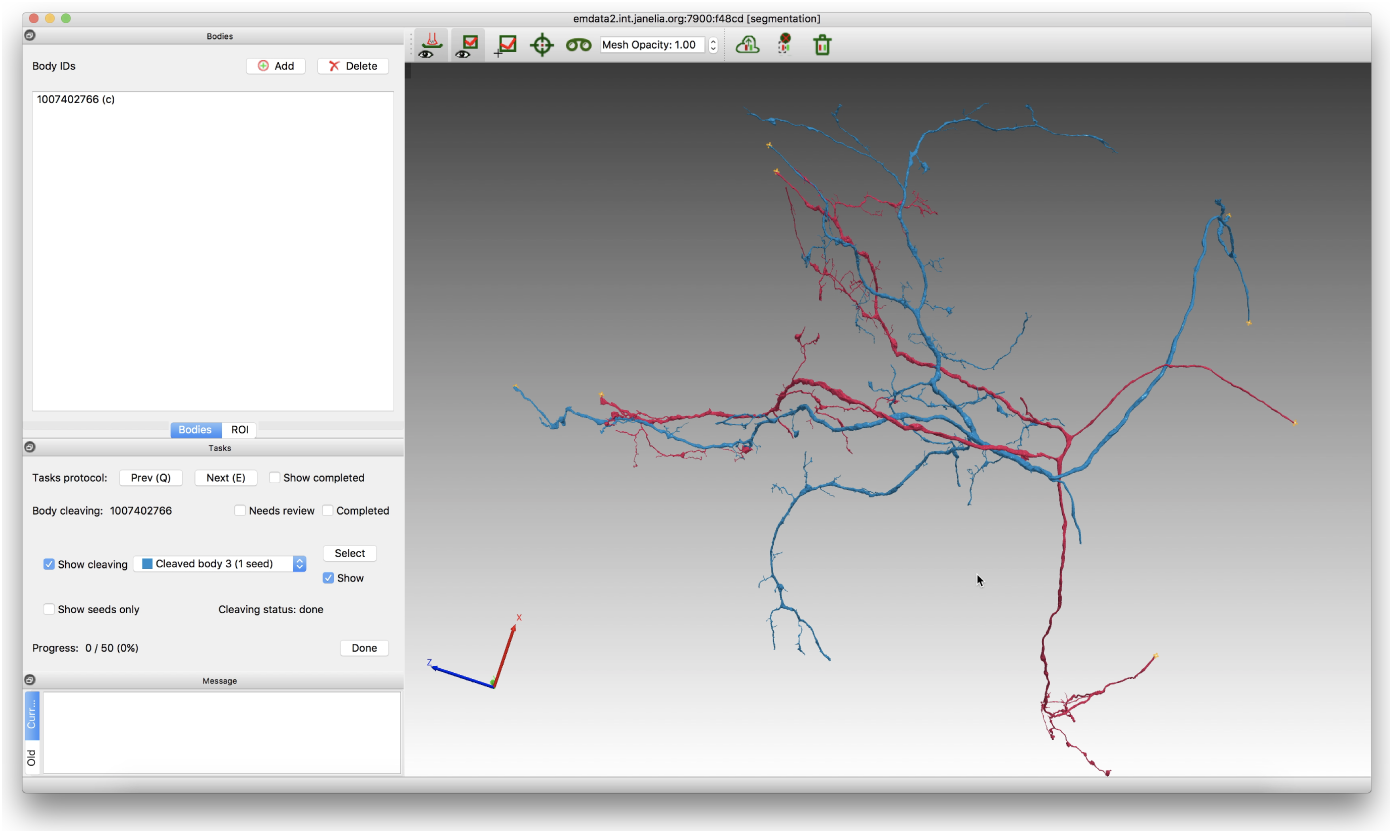
The Neu3 user interface for cleaving, showing the final two bodies.

**Figure 8:**
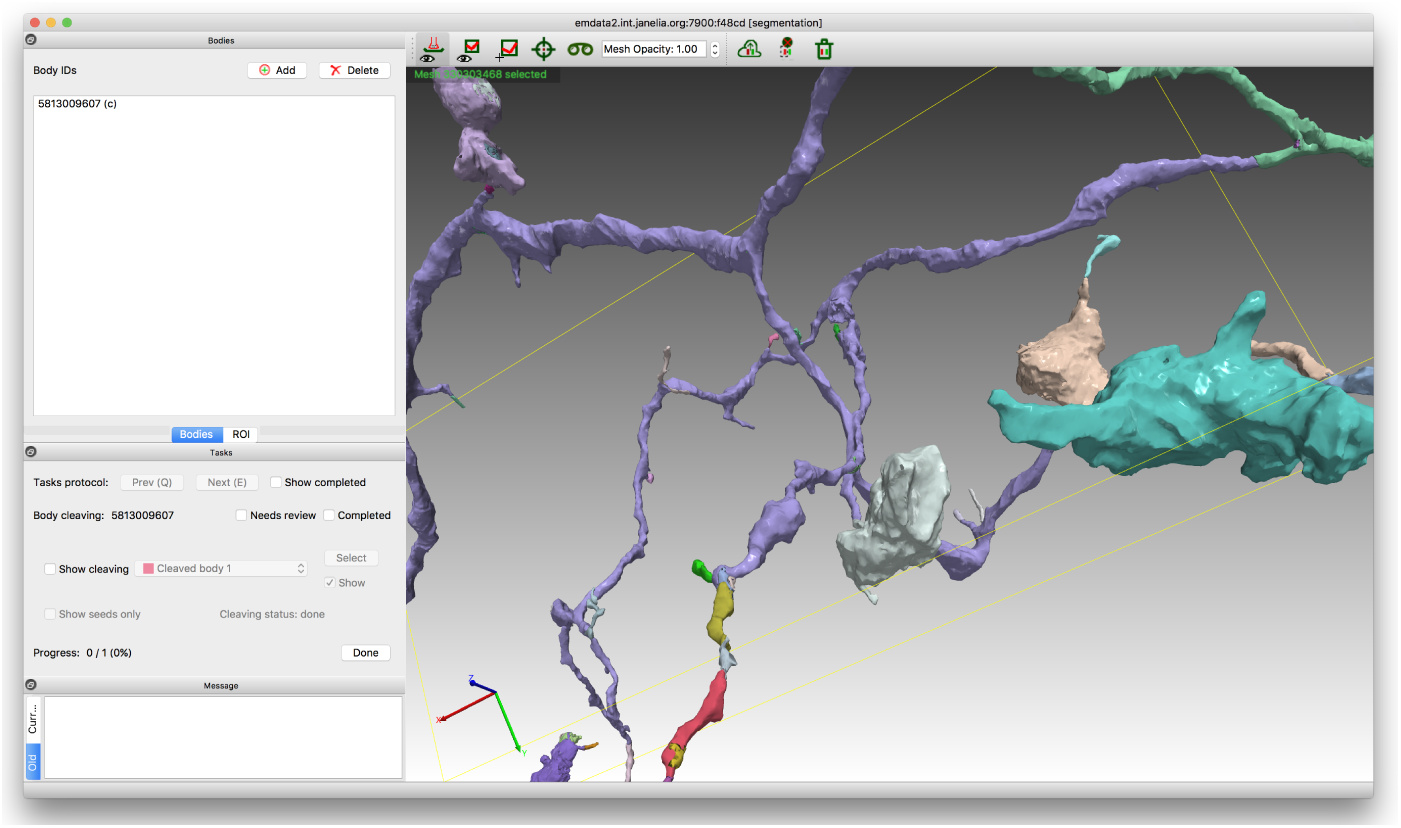
The Neu3 user interface showing a supervoxel, in purple, containing a false merge and needing to be split.

**Figure 9:**
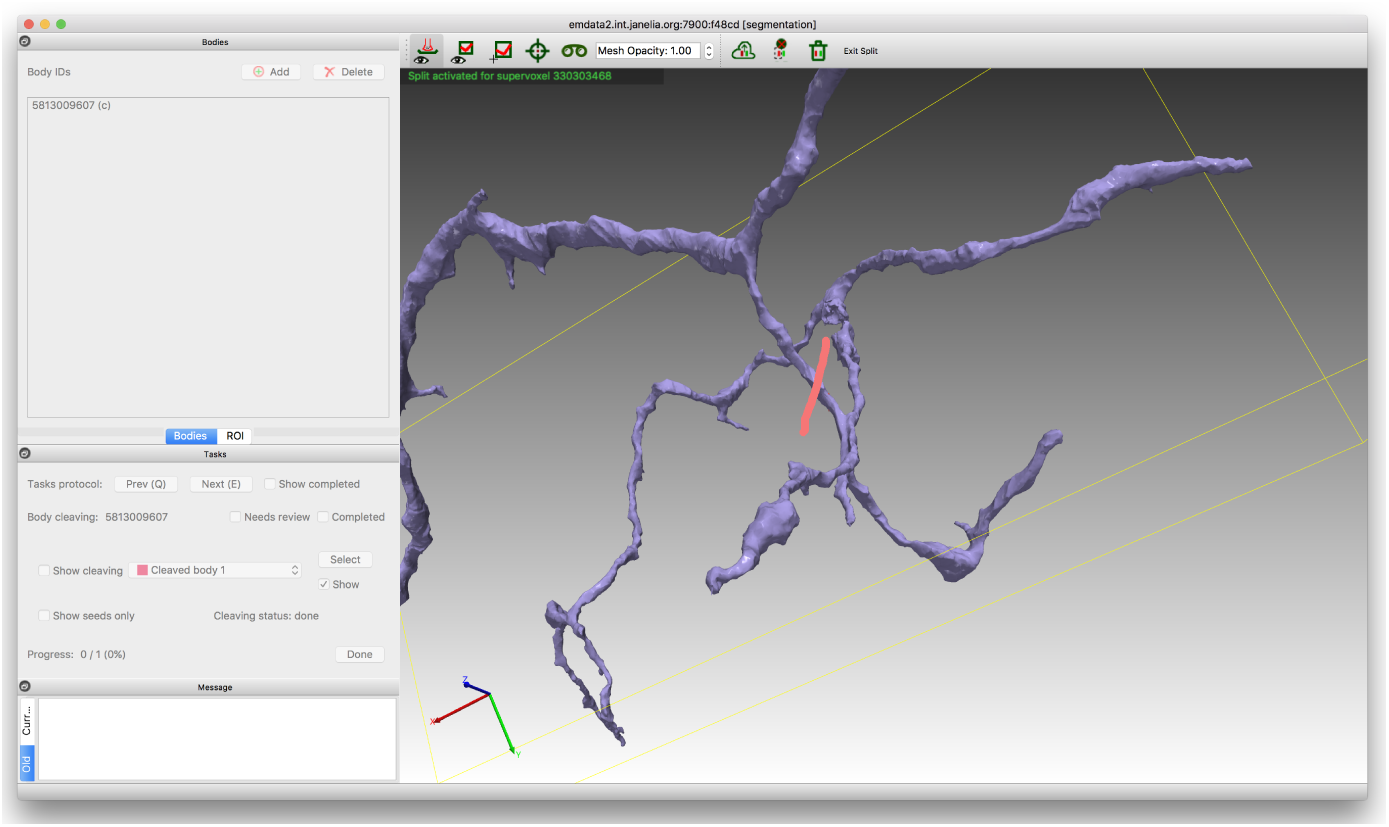
The Neu3 user interface for supervoxel splitting. The proofreader has isolated the supervoxel and painted a red stroke to seed the first split body.

**Figure 10:**
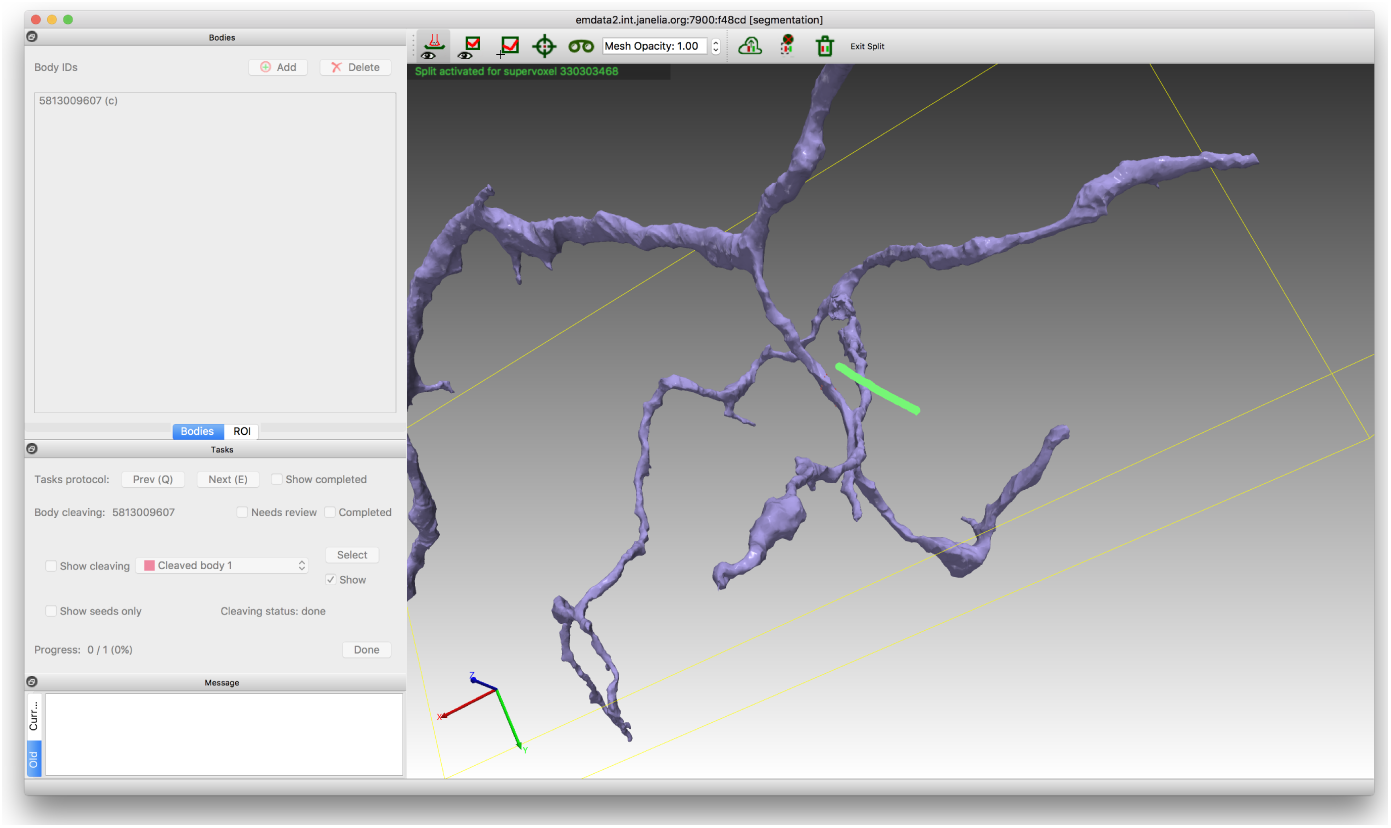
The Neu3 user interface for supervoxel splitting. The proofreader has painted a green stroke to seed the second split body.

**Figure 11:**
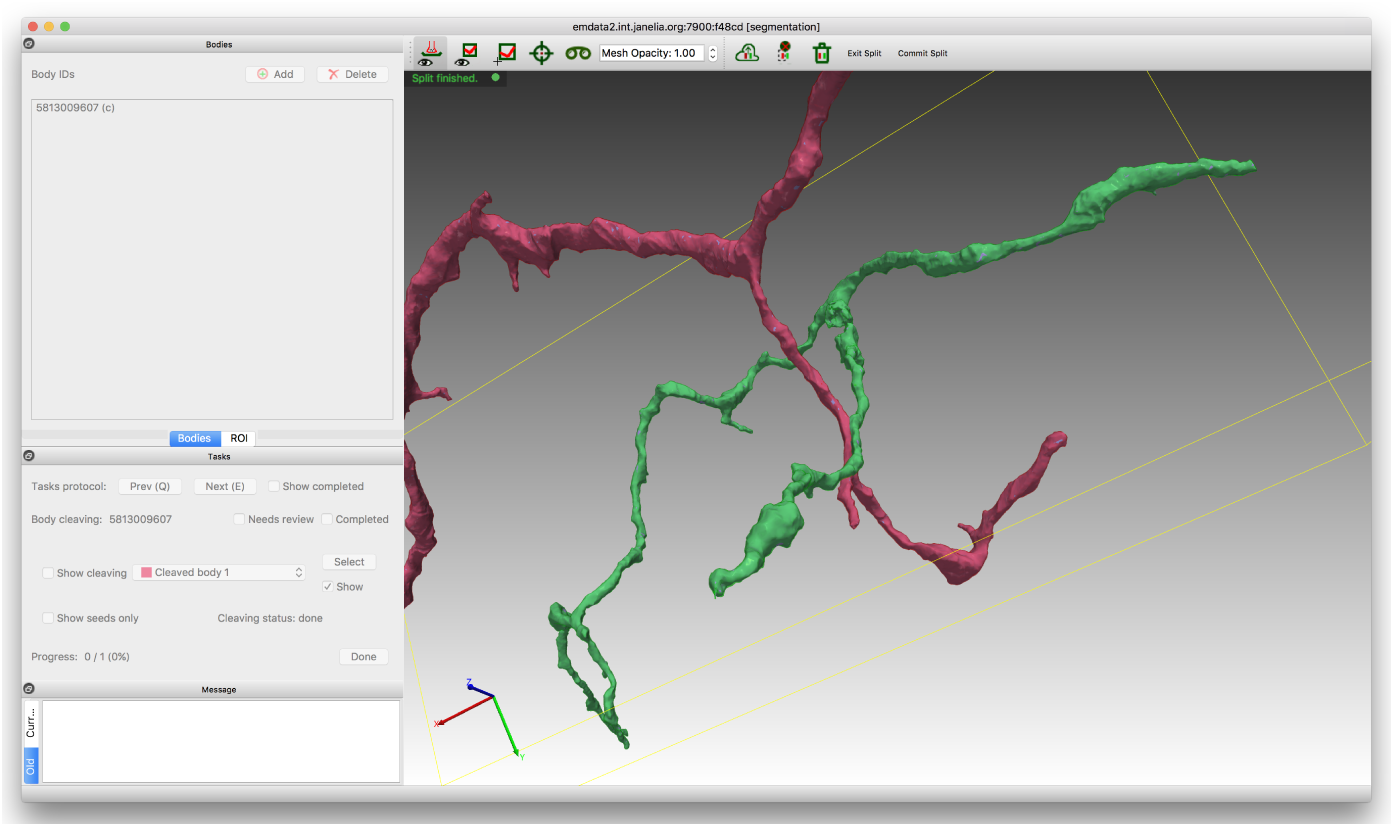
The Neu3 user interface, showing the result of supervoxel splitting.

**Figure 12:**
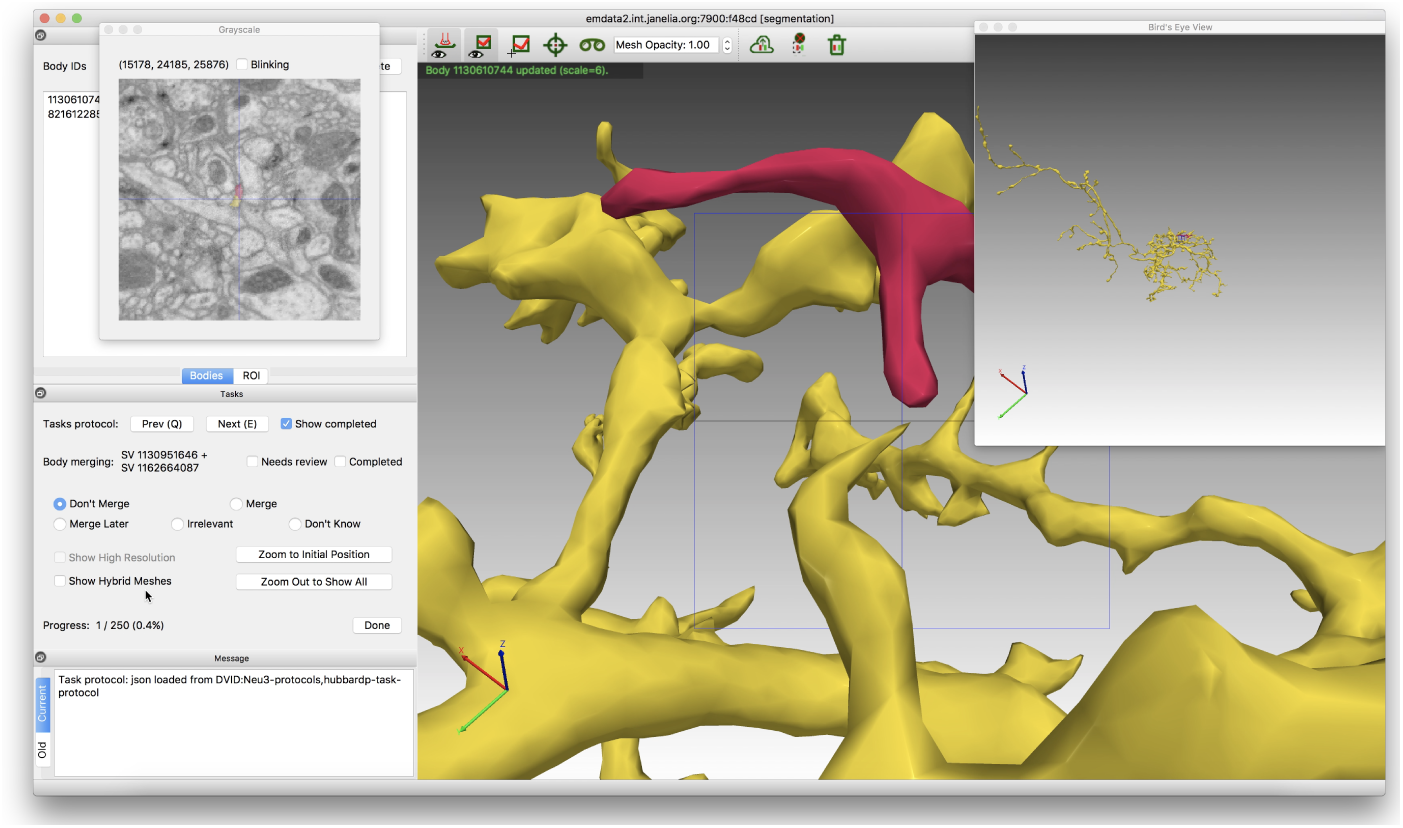
The Neu3 user interface, showing focused merging without hybrid visualization. It is not clear whether the gap at the cross-hairs is a true separation or an artifact of the low-resolution meshes.

**Figure 13:**
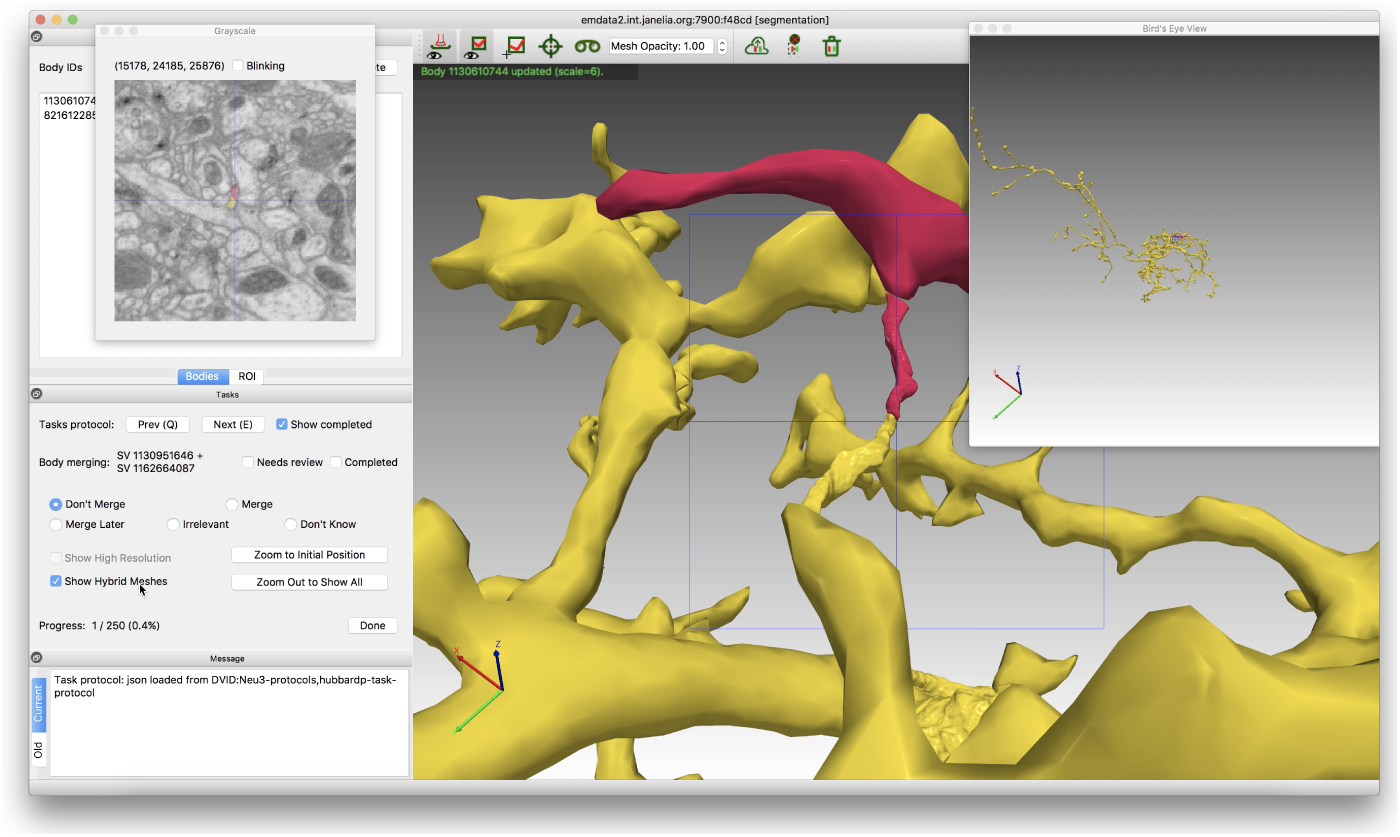
The Neu3 user interface, showing focused merging with hybrid visualization. Now the gap at the cross-hairs has disappeared and it is clear the bodies should be merged.

The camera for rendering the hybrid visualization is oriented initially to look at the merge position, with its view direction perpendicular to the line segment, *L*, and its “up” direction remaining as most recently set by the user. The camera is translated along the view direction so the smaller of the two bodies just fits in the rendered view.

Focused merging includes two additional views to give the proofreader more context. The first displays the grayscale data in the ROI with coloring for the task’s two bodies, to show features outside the two bodies. This 2D image comes from data resampled on the 3D plane through the merge position and normal to the view direction. The second additional view is a bird’s-eye view, to show the full morphology of both bodies in the task as a form of global context. It uses a camera oriented like the main camera, but translated out until both bodies just fit in the rendered view. Focused merging also allows the proofreader to disable hybrid visualization in favor of loading the precomputed, high-resolution supervoxel meshes for the two bodies if that (typically slower) approach is preferred.

For supervoxel splitting, the proofreader does not describe the exact splitting boundary, but instead paints strokes on the supervoxel’s mesh that lie within the distinct split regions. From these strokes, Neu3 infers seeds for a seeded watershed algorithm that performs the splitting. Neu3’s algorithm for inferring seeds extends the approach in NeuTu [6] to work on 3D meshes. Points on the skeleton of a stroke define the starting point for rays through the mesh in the camera view direction, and for each ray the segment between the first mesh entry and exit points is uniformly sampled to define seed points. The Neu3 splitting algorithm, run locally, is a further optimized, multiscale version of the offline NeuTu algorithm.

## 4 Semi-Automated Re-Segmentation

### 4.1 Data Representation

Our segmentation volume is represented as a set of supervoxels and an agglomeration mapping, which designates how subsets of supervoxels are grouped together to comprise the neuron fragments (segments) of the volume.

By assuming that the supervoxels are each atomic (i.e. none of them are internally undersegmented), all changes to the segmentation can be performed using simple edits to the agglomeration mapping. Such edits are executed as a series of merging and cleaving operations, in which sets of supervoxels are grouped together under the same agglomeration label for merging, or are relabeled to have different agglomeration labels for cleaving. Once a human proofreader has located the site of an over- or under-segmentation error in the agglomeration and specified a revised supervoxel grouping, applying the necessary merge/cleave operation to the agglomeration mapping is nearly instantaneous. But finding such errors specifying the revised grouping without automated assistance is time-consuming.

To facilitate rapid semi-automated editing, we precompute a region adjacency graph (RAG) of supervoxel-to-supervoxel adjacencies, stored as a table. Each row in the table lists a pair of supervoxels which are spatially adjacent, and an affinity score, indicating the likelihood that the two supervoxels belong together in the same agglomerated segment. The agglomeration labels are not stored in the graph.

### 4.2 Focused Merging

As described in Section 2, focused merging involves a pair of bodies that are candidates for merging. The candidates can be automatically preselected using various methods. Our method is to simply select rows from the RAG which have favorable affinity scores but have not already been merged together in the current version of the segmentation. Alternative methods may be based on morphological correlations between neighboring supervoxels, or be based on comparisons with other automated segmentation methods. Once a pair of objects has been selected as a candidate for merging, we precompute a focal point for the proofreading viewer. If the two candidate bodies are directly adjacent (actually touching), then we choose a point near the centroid of their shared border. If the objects do not quite touch due to insignificant classification errors, we find the points at which the objects come closest to each other, and choose the midpoint between them as the focal point.

### 4.3 Cleaving

When cleaving, as described in Section 2, the proofreader marks a sparse subset of supervoxels as seeds belonging to distinct bodies, and the neighboring un-marked supervoxels are automatically assigned labels according to their transitive affinities to the seeds. The label assignment is computed by a *cleave server* running on a separate machine, which performs a graph cut on the relevant subset of nodes and affinities from the RAG. The exact graph cut algorithm it uses is configurable. In practice, we use Kruskal’s algorithm [12] to find a minimum spanning forest among the supervoxels within the body of interest, seeded from the user’s annotations. More sophisticated alternatives such as hierarchical agglomerative clustering did not yield significantly better results, and were slower to compute, hindering interactive responsiveness.

#### 4.3.1 Cleave Server Implementation

The cleave server exposes an HTTP API for the Neu3 client to access. It has access to the RAG, and to a DVID server in which the segmentation is stored. The RAG itself is largely immutable. When a client wishes to perform a cleave computation for a particular object, the client’s request to the cleave server includes the object label, the marked supervoxel labels, and the DVID address the client is working with. To determine which subset of the RAG to use, the server fetches from DVID the list of component supervoxels that comprise the user’s object, along with a unique *mutation identifier* for the exact version of that object.

The server does not require the RAG to contain all possible supervoxel adjacencies in the volume. There are multiple reasons the RAG might be missing an adjacency entry for a pair of nearby supervoxels. For example, if the initial segmentation left a gap between two supervoxels that do, in fact, belong to the same object, the RAG construction algorithm might not have considered them adjacent, even though a human would. Another example occurs when a supervoxel is split into multiple supervoxels after the RAG was initially generated. In cases where intra-object supervoxel adjacencies are missing from the RAG, it is possible that the subgraph for the object’s supervoxels will constitute more than one connected component, rather than a single connected graph. In those cases, the cleave server fetches segmentation data from DVID and analyzes it to insert additional edges into the subgraph based on spatial proximities, until all connected components can be joined. Once the necessary edges have been obtained, the cleave operation can be performed on the now fully connected subgraph. In addition, the subgraph is cached along with the object’s mutation identifier, so that subsequent client requests for the object can be serviced without re-fetching the supervoxel list from DVID and reconstructing the subgraph from scratch. The cached subgraph remains valid until the object is modified in DVID, in which case the cleave server detects that the object’s mutation identifier has been changed, and the previously cached subgraph can no longer be used.

Note that the cleave server does not send cleave commands to DVID. Instead, the results of the cleave operation (i.e., a proposed partitioning of the object’s supervoxels) is returned to the client, whose user may or may not choose to commit the new partitioning to DVID via a cleave command.

## 5 Proofreading Workflow Details

The Neu3 application used for proofreading is a desktop application, implemented in C++ using Qt for the user interface and OpenGL for rendering. Most proofreaders run Neu3 on Linux, although it does run on macOS. Neu3 provides general user-interface controls common in 3D applications, like mouse-based camera manipulation and mesh selection. It also features a section in the lower-left corner of the main window for controls particular to the protocol in progress, as described in Sections 5.1, 5.2, and 5.3. Keyboard shortcuts are the proofreaders’ preferred way of controlling the protocols, allowing one hand to keep the mouse cursor in the 3D view for efficiency. Thus we put relatively little effort into optimizing the arrangement of buttons and other widgets.

### 5.1 Cleaving

A cleaving task starts by displaying all the supervoxels of an agglomerated body, with all supervoxel meshes having the same color. Figure 3 shows an example of a body that contains a false merge and needs to be cleaved.

A proofreader can toggle the rendering to color each supervoxel independently, as shown in Figure 4. Doing so can give insight to where there might be distinct bodies.

To begin cleaving, the proofreader toggles back to the original coloring mode, and clicks to select a mesh that belongs in the first body. Pressing the spacebar designates this supervoxel as a seed for the first body, and changes the color of the mesh to red, the color for the first body, as shown in Figure 5. Neu3 sends this seed in an asynchronous request to the cleave server, and the server sends back a reply indicating how the supervoxels are partitioned among the bodies. With one seed, all supervoxels belong to the same body.

The proofreader then presses a number key to indicate that subsequent seeds will belong to a different body. Selecting another mesh and pressing the spacebar sets such a seed, and gives the mesh another body color, as shown in Figure 6. Neu3 sends both seeds to the cleave server, and gets back an updated assignment of supervoxels to bodies, as shown in Figure 7.

In this example, the proofreader has chosen seeds wisely, and the two seeds are sufficient to fix the false merge in the original body. If the results are not satisfactory, the proofreader can undo the setting of seeds and set additional seeds for up to twenty bodies.

When satisfied with the results, the proofreader marks this task as completed. Neu3 uses DVID’s REST API to perform the cleaving operation on the stored segmentation, cleaving supervoxels off the original body. To avoid conflicts with other proofreaders, Neu3 uses a simple librarian service to keep the original body “checked out” and inaccessible to other proofreaders for the duration of this task. Once the task is completed, Neu3 “checks in” the body and proceeds to the next task assigned to the proofreader.

### 5.2 Supervoxel Splitting

Supervoxels containing false merges are rare, but if encountered during a cleaving task, such a supervoxel can be split immediately. Figure 8 shows an example. The proofreader selects the supervoxel mesh and uses a context menu to enter the supervoxel split mode.

The proofreader then paints one or more strokes that overlap the part of the supervoxel that belong in the first split body. Before doing so, the proofreader can hide all other supervoxels to reduce distractions. See Figure 9.

After pressing a key to switch to the next split body, the proofreader paints another stroke to define this body’s seeds. Figure 10 shows this stroke.

This supervoxel appears to involve only two falsely merged strands, so the seeds for two bodies should be sufficient. The proofreader then presses shift-spacebar to trigger the splitting algorithm. When the algorithm is finished, Neu3 displays the two new bodies, as seen in Figure 11. The split bodies look correct, so the proofreader can commit the split. The split supervoxel is replaced with two new ones, and their meshes are generated. The proofreader then can redisplay all the other supervoxels and continue cleaving as described above.

A proofreader who prefers not to address the false merge immediately can instead use Neu3’s support for “todo” annotations. In a special mode, the proofreader can click at a location on a body and choose from several types of annotations, which Neu3 stores in DVID with the 3D position corresponding to the click. These annotations are displayed as special marks in the 3D view. The annotation records in DVID can be mined periodically to generate new tasks for proofreaders to perform.

### 5.3 Focused Merging

Each focused merging task is meant to be simpler and faster to perform than a cleaving task, since we expect the number of focused merging tasks to be greater.

The proofreader simply indicates whether the two bodies in the tasks should be merged or not. The user interface does present a few other choices, which are less commonly needed. “Merge Later” is appropriate if the proofreader decides that one of the bodies needs further cleaving. “Irrelevant” is appropriate if the proofreader determines that one of the bodies is a glial cell that was incorrectly categorized in the initial segmentation. “Don’t Know” indicates a case that is too ambiguous for one proofreader to decide, and requires more discussion.

The proofreader can choose not to use the hybrid visualization discussed in Section 3. Omitting the higher resolution the ROI can lead to ambiguity, though, as in Figure 12, where it is not clear if the two bodies actually touch. It helps to consult focused merging’s two additional views, the grayscale view of the original volume data and the bird’s-eye view of the two bodies, but leaving hybrid visualization on can be most helpful; see Figure 13.

In this example, hybrid visualization shows the bodies should be merged, and this decision is further confirmed by the grayscale view. The proofreader thus chooses the “Merge” option. Figure 14 shows how the mesh coloring is updated to indicate that the two bodies are now one.

**Figure 14:**
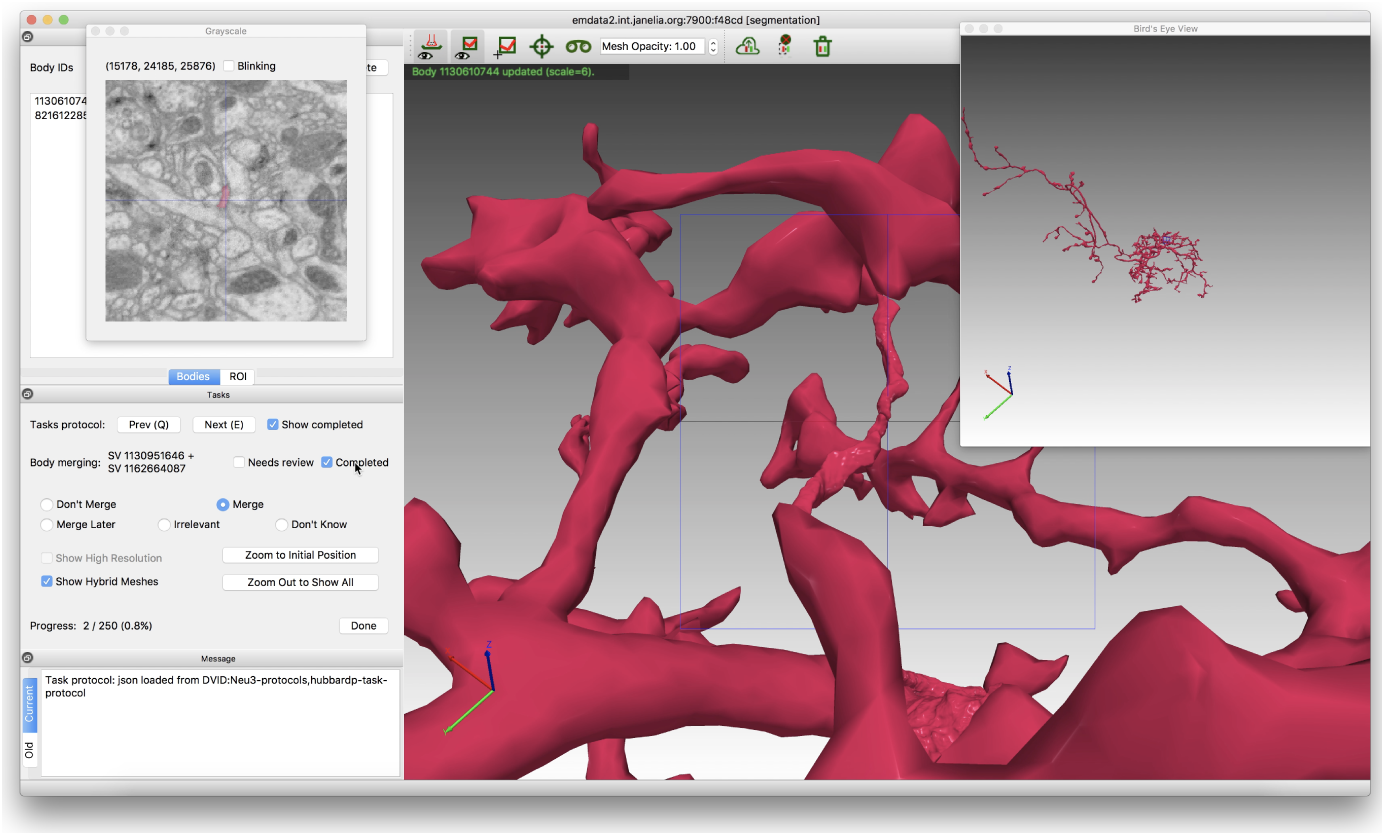
The Neu3 user interface, showing the choice to mege in a focused merging task.

The proofreader now marks the tasks as completed. For historical reasons, Neu3 does not directly update the segmentation stored in DVID as it does for cleaving tasks. Instead, it stores information about the proofreader’s decision in an ad hoc DVID data instance, which is processed later by scripts that make the changes to DVID’s segmentation. This delayed approach allows for additional quality control based on biological constraints, which is more important for focused merging since there is some risk of merges that make sense independently combining into a whole that does not make sense. Nevertheless, updating Neu3 to directly update the DVID segmentation is on the agenda for future work.

## 6 Results

### 6.1 Cleaving

We evaluated cleaving with a user study. From an unpublished dataset, we chose 20 bodies with one significant false merge, and 22 with no significant false merge. The bodies are large but not huge, coming from the second thousand bodies in the segmentation when ordered by volume. The participants in the test were seven proofreaders with extensive experience using cleaving in production. The instructions for the test were to find each task’s most significant false merge, if one exists.

Agreement between proofreaders in the test is measured in terms of synapses, since proper assignment of synapses to bodies is among the most important criteria for a connectome. Specifically, each cleaving task can be thought of as set of categorical decisions: for each synapse, does it belong with the original body, or with a cleaved-off body? Our proofreading convention is that when cleaving into multiple pieces, the largest piece should retain the identity of the original body. In this test, there were some tasks where the two pieces were roughly the same size, and it was arbitrary which piece was designated as the original body. To prevent this arbitrary choice from appearing as false disagreement, we rephrased the categorical decisions slightly: for each synapse, does it belong with the majority body (having the majority of the synapses) or the other body? To compare the agreement of multiple parties making such categorical decisions, a common statistic is Fleiss’s kappa [13]. The kappa value ranges from -1 to 1, with values closer to 1 indicating agreement better than would be expected by chance, and 1 indicating perfect agreement. Figure 15 presents the kappa values from our tests, with the bars in relative order of increasing task complexity, measured as the number of branches in a skeleton for the task’s body. Cross-hatched bars indicate cases of perfect agreement where Fleiss’ kappa is undefined (its denominator is zero when all proofreaders assign all synapses to the same body); all tasks with no false merge are such cases. Overall, the agreement rates are high. The kappa values below 1 reflect that three bodies with a false merge (7% of all the bodies) were not cleaved by one of the proofreaders, and one body with a false merge (2%) was not cleaved by two of the proofreaders.

**Figure 15:**
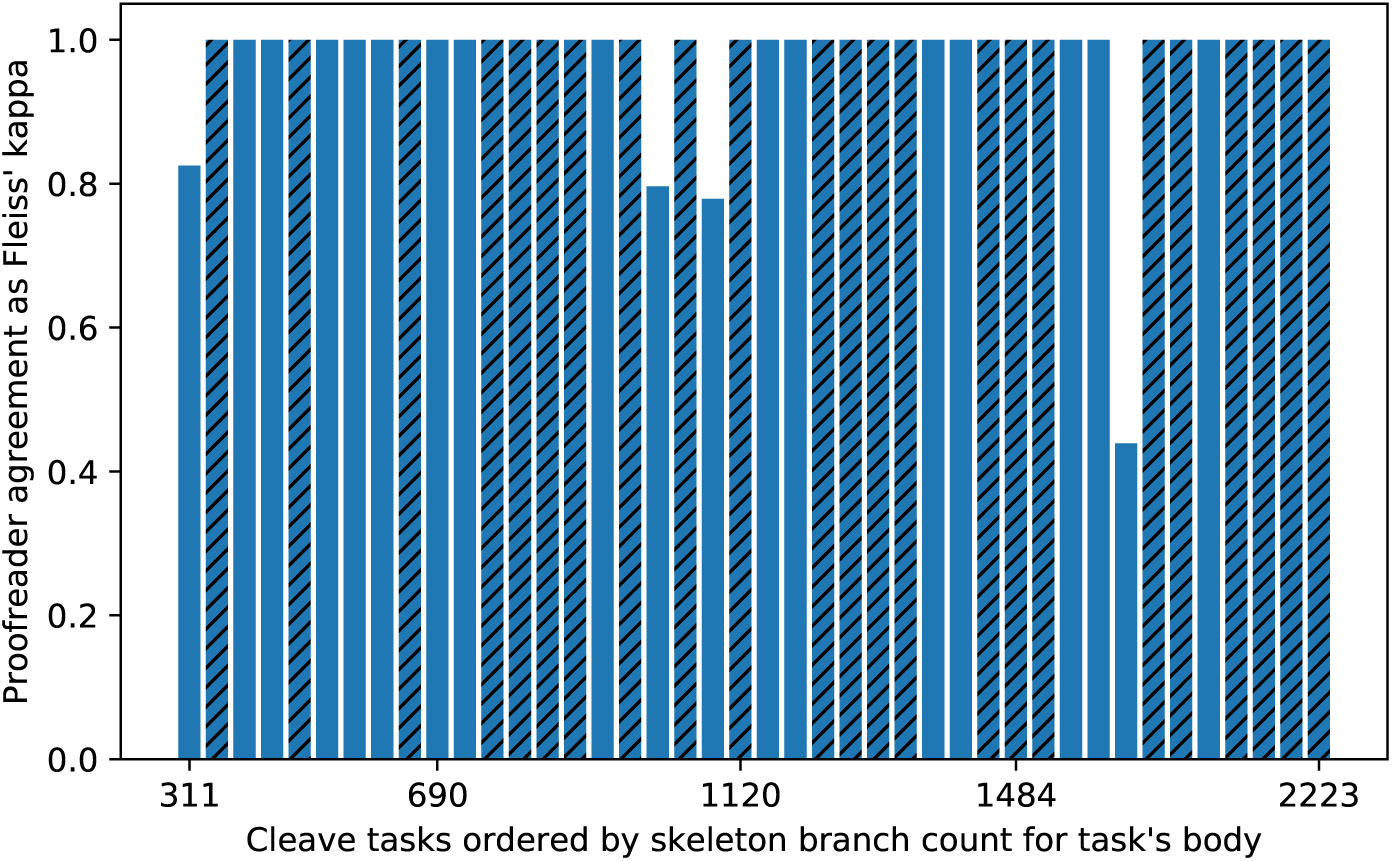
Proofreader agreement for a cleaving test, as Fleiss’ kappa, with values closer to 1 indicating more agreement, beyond what would be expected by chance. A task’s kappa compares how each synapse from the task’s body was assigned to cleaved bodies by each proofreader. Cross-hatching indicates cases of perfect agreement where Fleiss’ kappa is undefined (e.g., there was no false merge and no proofreader did any cleaving).

Performance assessment of the cleaving test yields the main conclusion that cleaving scales well. Figures 16, 17 and 18 show that the mean time spent on each cleaving task grew only modestly with the complexity of the task, using three possible measures of complexity: the number of branches in the skeleton form of the task’s body, the number of supervoxels in that body, and the number of voxels in that body.

**Figure 16:**
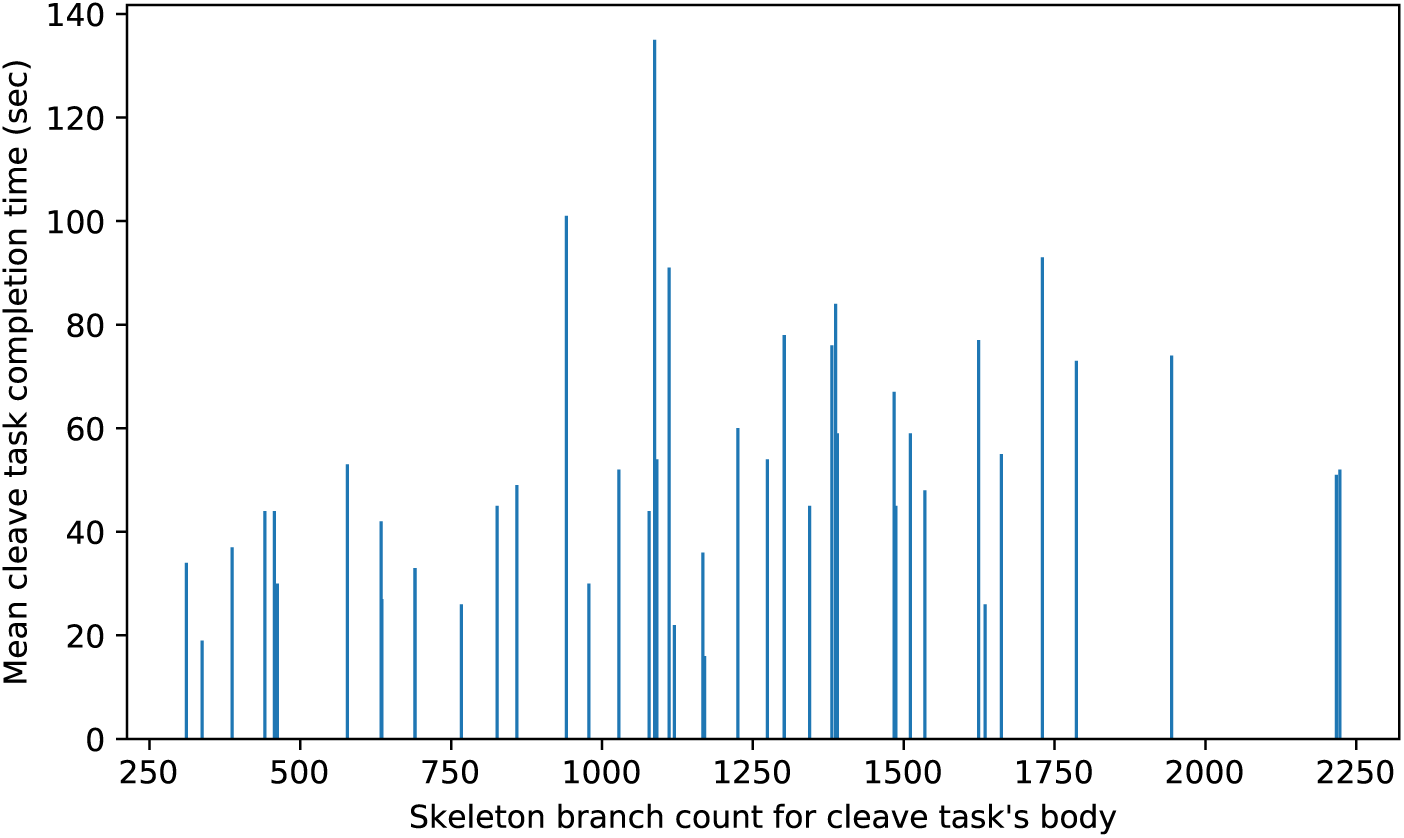
Mean proofreader time for cleaving tasks in a test grows only modestly with body complexity, measured as skeleton branch count.

**Figure 17:**
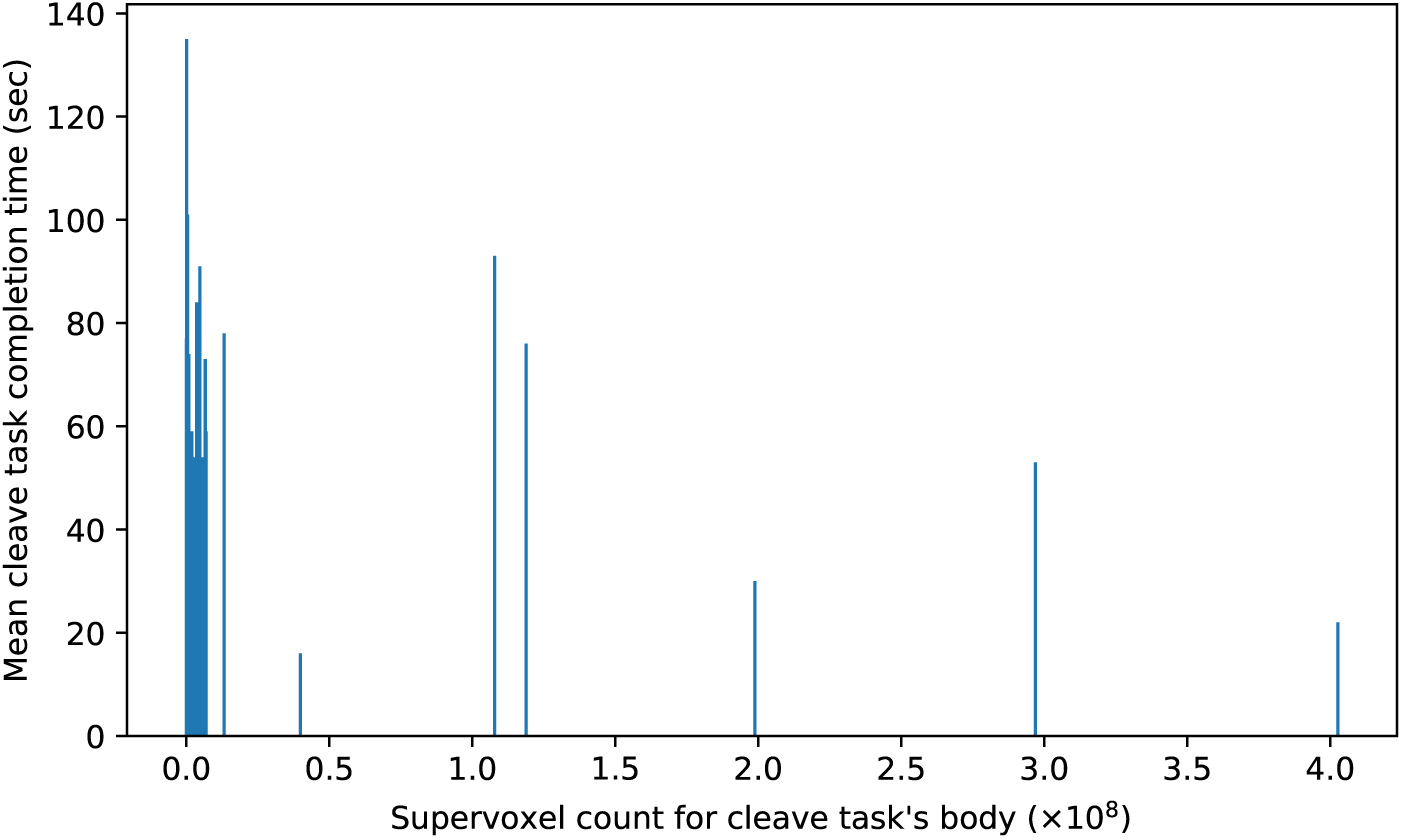
Mean proofreader time for cleaving tasks in a test grows only modestly with body complexity, measured as supervoxel count.

**Figure 18:**
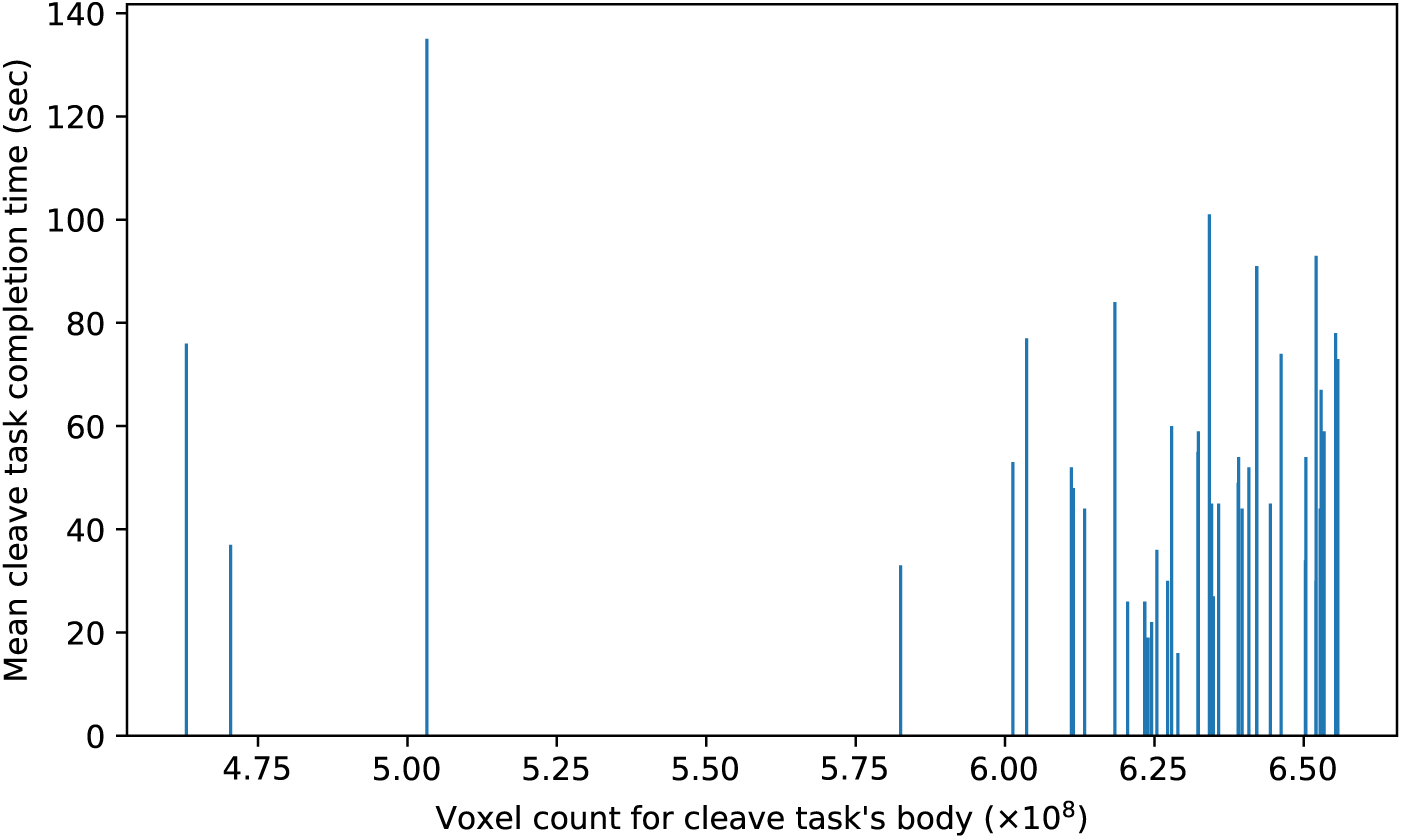
Mean proofreader time for cleaving tasks in a test grows only modestly with body complexity, measured as voxel count.

There was some variability in the time per cleaving task. Figure 19 shows box plots of the time per tasks, with tasks ordered by increasing complexity (skeleton branch count). Variability, indicated by the range between quartiles, appears to increase slightly with task complexity. Figure 20 shows box plots for only the tasks with one false merge, and Figure 21 shows only the tasks with no false merge. Variability appears slightly greater in the case of one false merge. A possible explanation for the variability is that some proofreaders are more meticulous and work more slowly, as shown Figure 22, which plots total time spent on all tasks per proofreader.

**Figure 19:**
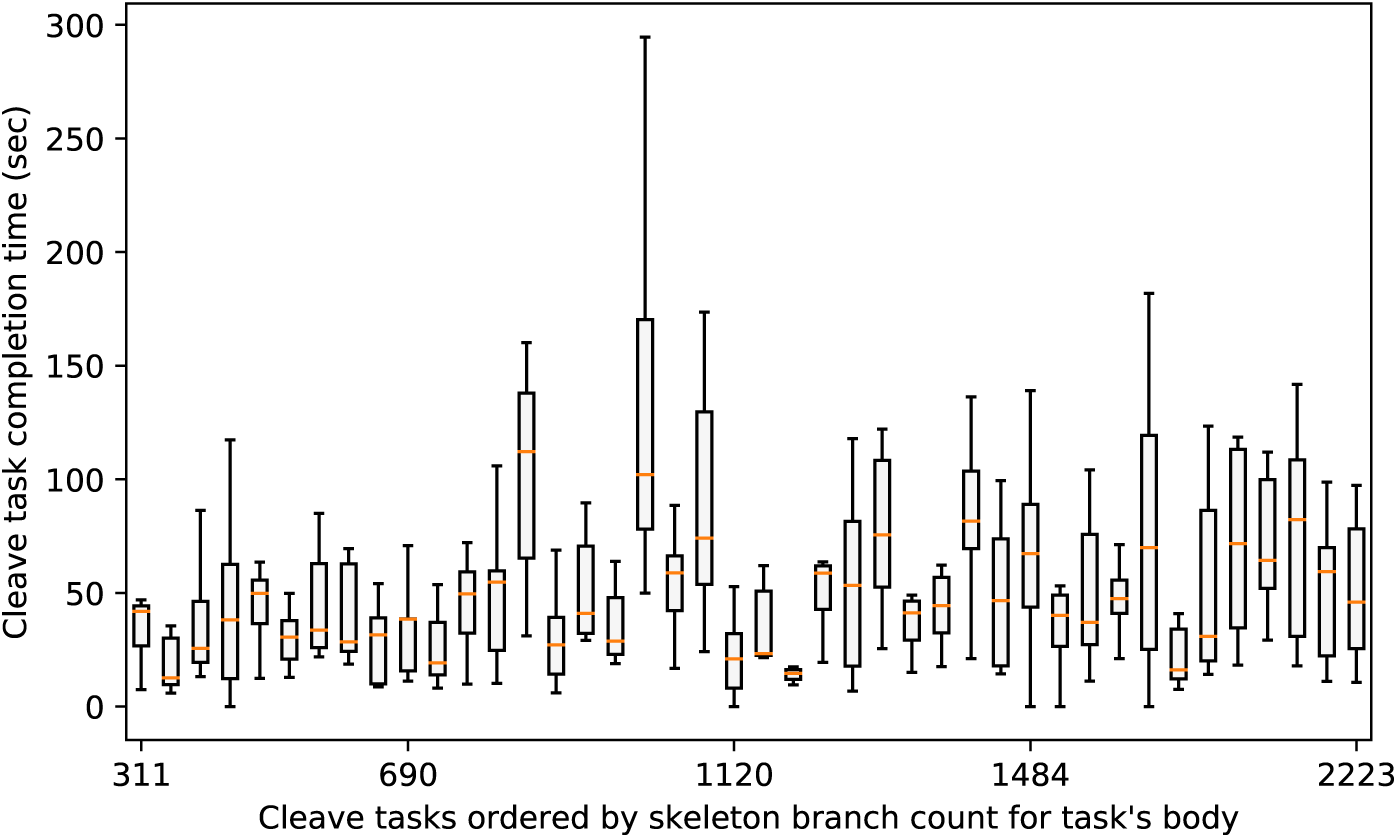
Cleaving task completion times as box plots (first quartile, median, third quartile, whiskers at 1.5 × IQR) show some variability, which increases slightly with task complexity.

**Figure 20:**
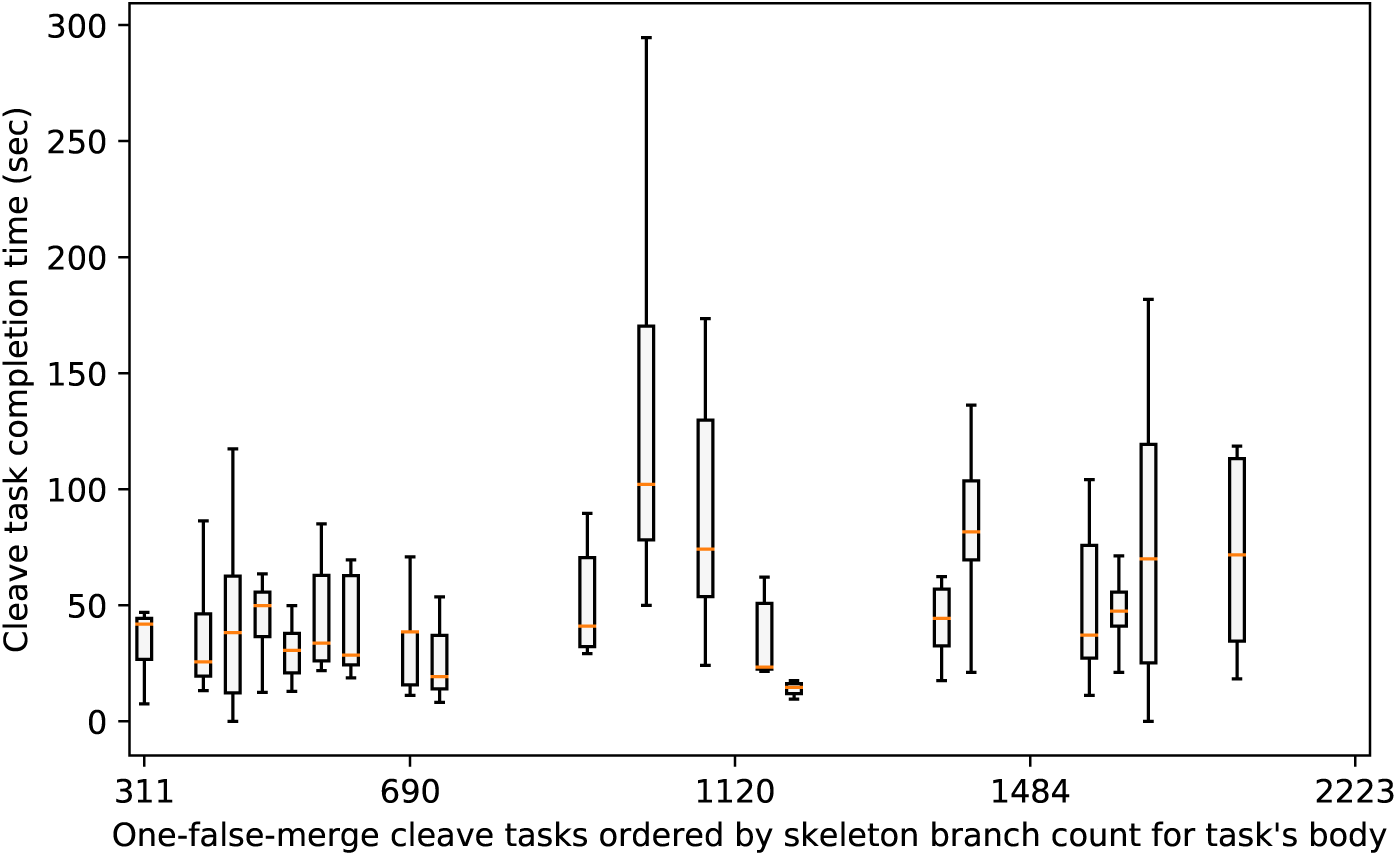
Completion times for cleaving tasks with one false merge, as box plots (first quartile, median, third quartile, whiskers at 1.5 × IQR).

**Figure 21:**
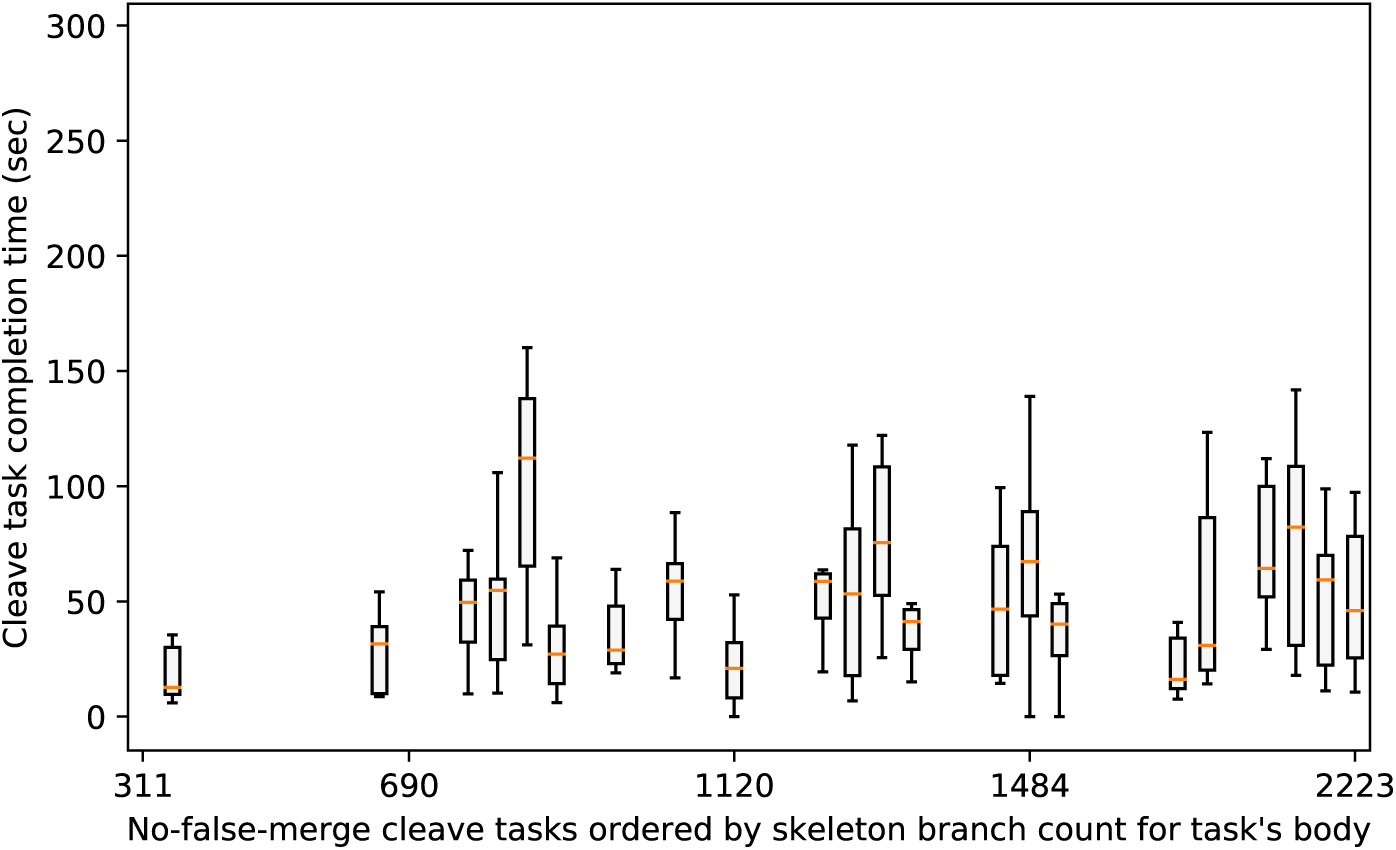
Completion times for cleaving tasks with no false merge, as box plots (first quartile, median, third quartile, whiskers at 1.5 × IQR).

**Figure 22:**
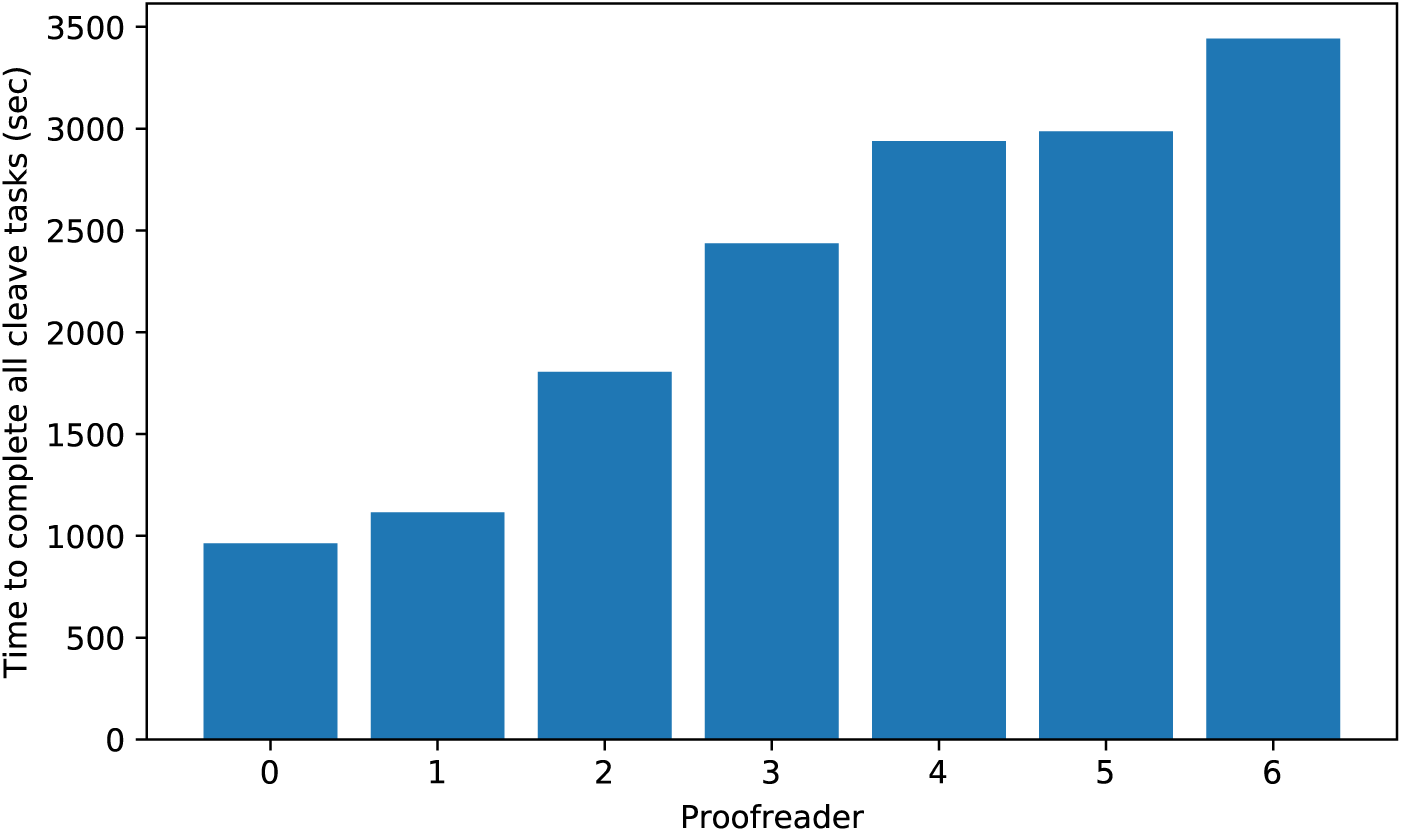
The total time for all cleave tasks varied across proofreaders in the test.

Another important conclusion from the tests is that cleaving feels interactive. Figure 23 shows box plots of the response time for the cleave server in the test. This time is the delay that the user would experience before seeing the results after placing a cleaving seed. The plots indicate that while there is some variability, response times and hence delays were consistently well below 0.5 seconds. This good performance was maintained even as task complexity grew.

**Figure 23:**
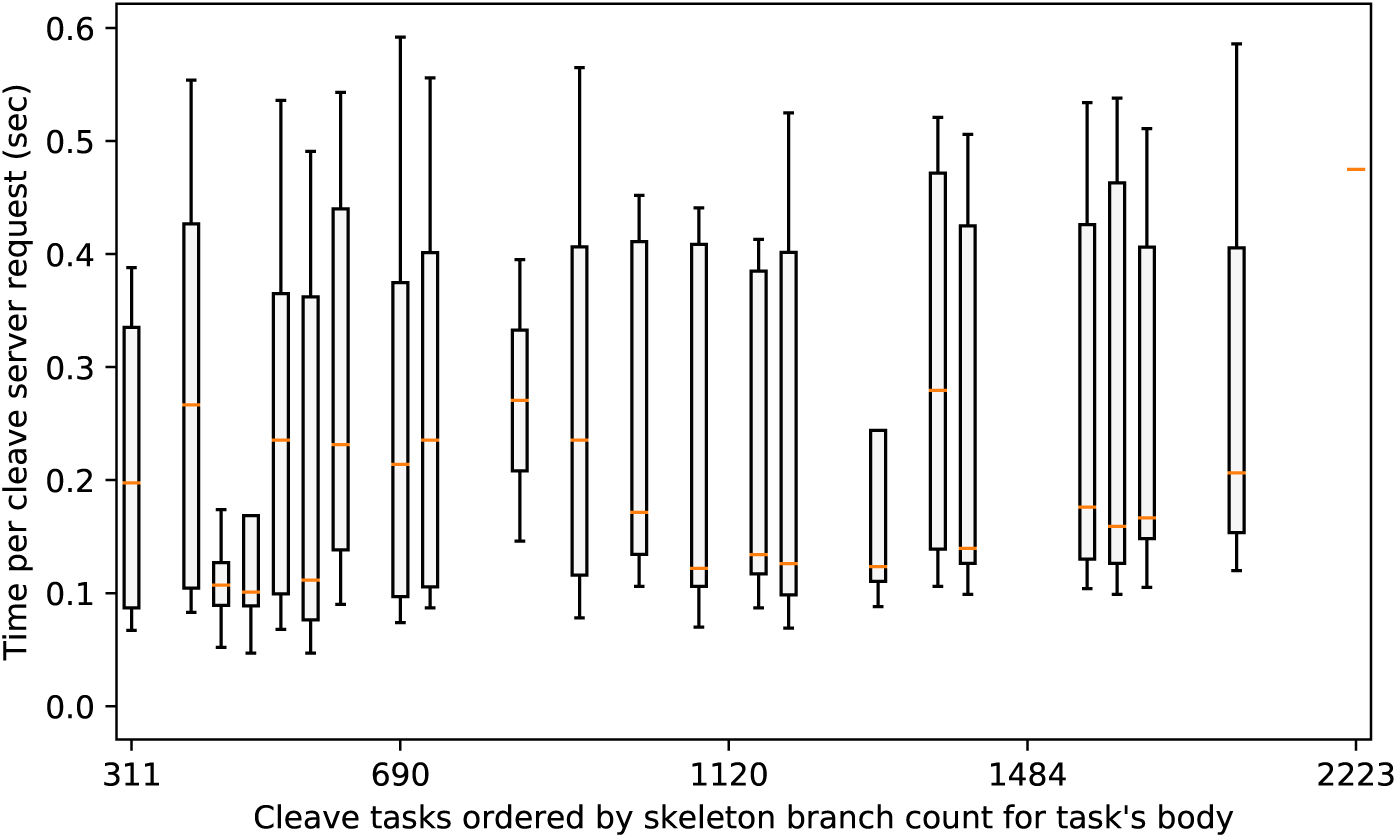
Box plots (first quartile, median, third quartile, whiskers at 1.5 × IQR) of the cleave server response for the test’s tasks indicate that response time was fast enough to feel interactive, even as task complexity grew.

### 6.2 Focused Merging

For focused merging, our evaluation was based on production work on an unpublished dataset. We evaluated proofreader agreement during a training phase. Twenty-two proofreaders performed five assignments of 50 tasks each to gain experience with focused merging. The assignments involved challenging data, with ambiguities in the grayscale and the segmentation. We compared the decision of each proofreader on each task to the decision from an experienced proofreader who helped to develop the protocol. This pairwise agreement rate was 95.78% over all 22 proofreaders. Based on the results of this training, one proofreader was assigned to tasks other than focused merging, so the pairwise agreement likely increased during production work.

Performance evaluation comes from a four month period of production work on that dataset. A total of just under 600, 000 decisions were made during this period. Figures 24 and 25 shows box plots of the task completion times, as an aggregate of all types of decisions, and broken down by decision type. Outliers, like proofreaders going home for the day in the middle a task, contributed to the spread in recorded times. Nevertheless, the median time overall was quite fast, around 7 seconds.

**Figure 24:**
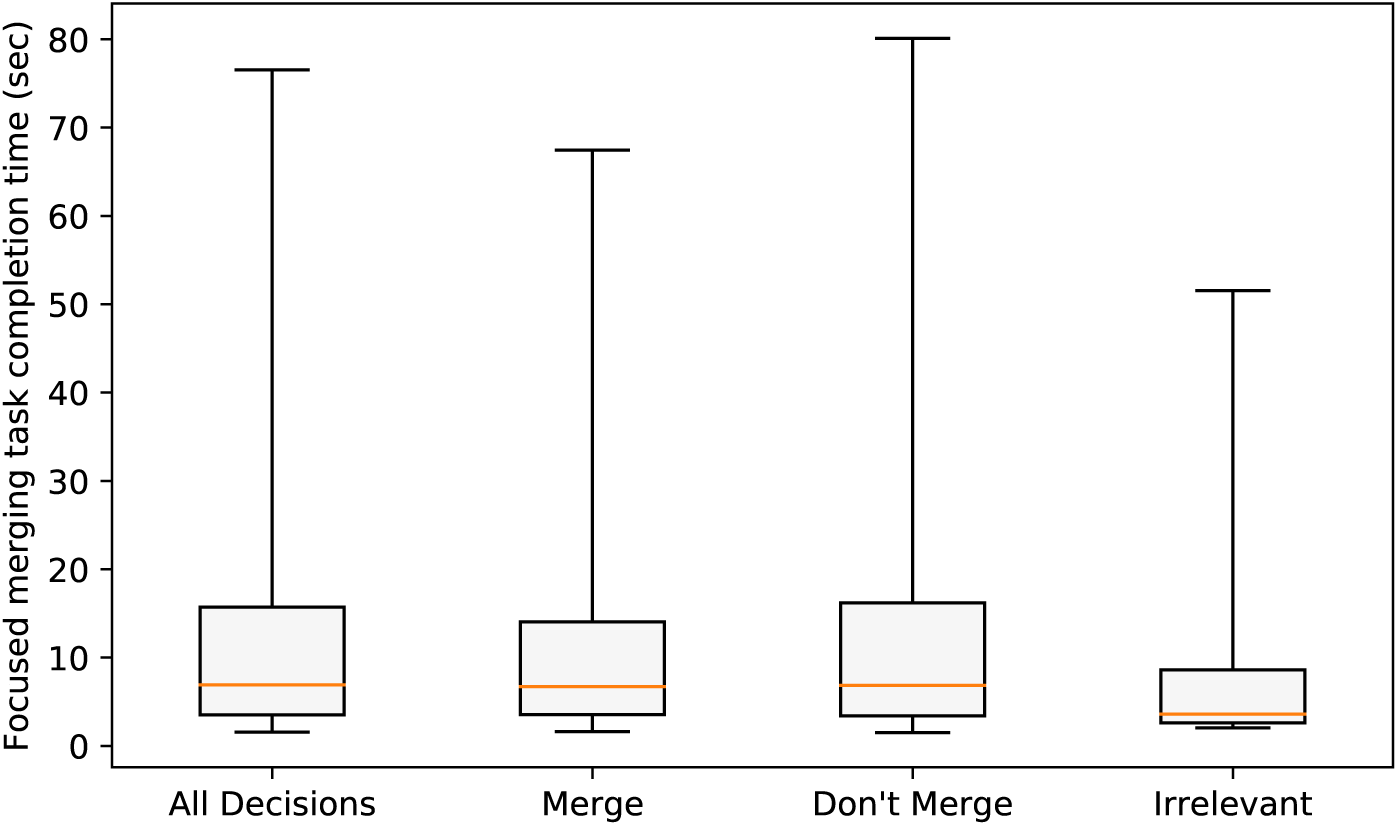
Completion times for focused merging tasks during production, as box plots (first quartile, median, third quartile, whiskers at 5th and 95th percentiles, due to outliers). The first box plot aggregates all types of decisions, and the others show specific types of decisions.

**Figure 25:**
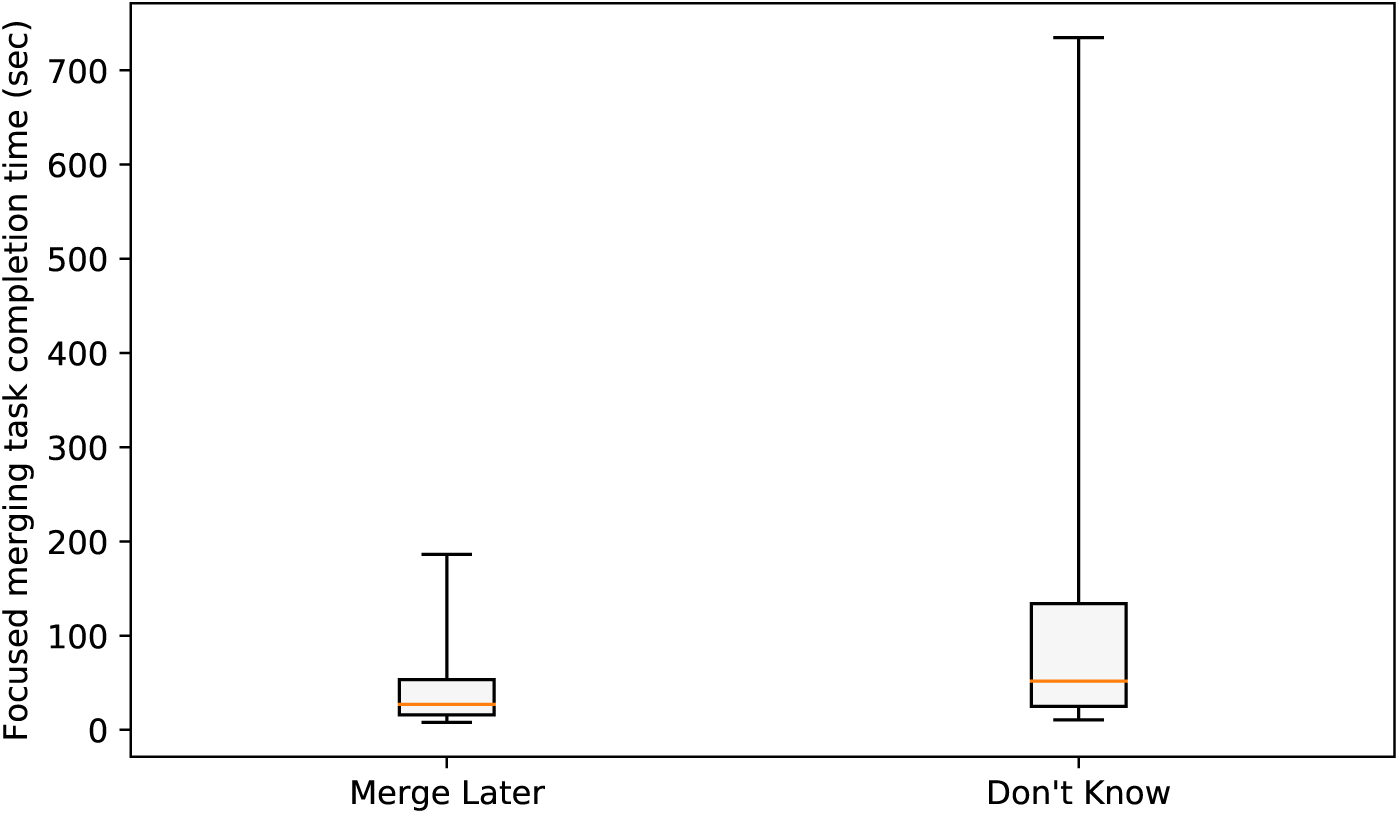
Completion times for focused merging tasks during production, as box plots (first quartile, median, third quartile, whiskers at 5th and 95th percentiles, due to outliers). These box plots show two types of decisions that took longer, and thus need a different scale.

### 6.3 Splitting

We measured the processing time for the splitting of 18 bodies, with various sizes ranging from 89 thousand to 90 million voxels. The processing time was roughly proportional to the number of voxels of the body, as shown in Figure 26. The average processing speed is about 1.4 million voxels per second, leading to significant waiting time for bodies with millions of voxels or larger sizes.

**Figure 26:**
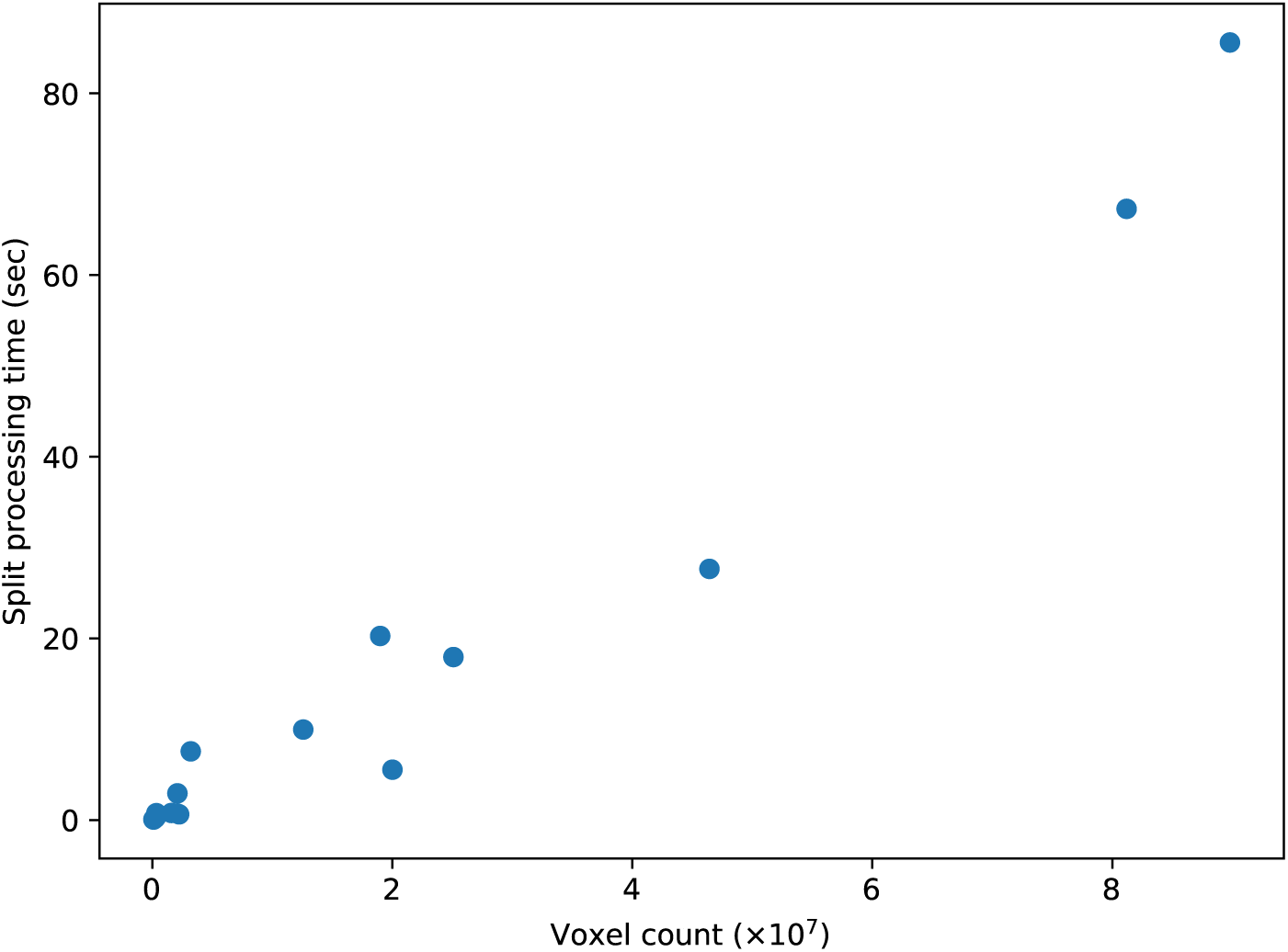
The processing time for splitting a body is roughly proportional to the size of the body.

## 7 Conclusions

The cleaving, supervoxel splitting, and focused merging techniques we introduced have been effective in producing a connectome for an unpublished dataset of unprecedented scale. To accomplish production at this scale, our tools emphasize high-throughput, top-down proofreading through streamlined protocols that first highlight the 3D morphology of the neurons. Most often, proofreaders can quickly make a decisions simply based on these 3D shapes. Furthermore, the ability to interactively cleave large neurons rapidly encourages a more aggressive approach to automatic segmentation and agglomeration. Since our workflow is more tolerant of large false merge errors, we can now tune agglomeration to have significantly fewer false splits.

## Acknowledgements

We would like to thank Christopher Knecht for advice on proofreader workflows, the proofreaders for their tireless work, and the entire FlyEM project team at the Janelia Research Campus for valuable discussions and support.

